# Phase-locking patterns underlying effective communication in exact firing rate models of neural networks

**DOI:** 10.1101/2021.08.13.456218

**Authors:** David Reyner-Parra, Gemma Huguet

## Abstract

Macroscopic oscillations in the brain have been observed to be involved in many cognitive tasks but their role is not completely understood. One of the suggested functions of the oscillations is to dynamically modulate communication between neural circuits. The Communication Through Coherence (CTC) theory establishes that oscillations reflect rhythmic changes in excitability of the neuronal populations. Thus, populations need to be properly phase-locked so that input volleys arrive at the peaks of excitability of the receiving population to communicate effectively. Here, we present a modeling study to explore synchronization between neuronal circuits connected with unidirectional projections. We consider an Excitatory-Inhibitory (E-I) network of quadratic integrate-and-fire neurons modeling a Pyramidal-Interneuronal Network Gamma (PING) rhythm. The network receives an external periodic input from either one or two sources, simulating the inputs from other oscillating neural groups. We use recently developed mean-field models which provide an exact description of the macroscopic activity of the spiking network. This low-dimensional mean field model allows us to use tools from bifurcation theory to identify the phase-locked states between the input and the target population as a function of the amplitude, frequency and coherence of the inputs. We identify the conditions for optimal phaselocking and selective communication. We find that inputs with high coherence can entrain the network for a wider range of frequencies. Besides, faster oscillatory inputs than the intrinsic network gamma cycle show more effective communication than inputs with similar frequency. Our analysis further shows that the entrainment of the network by inputs with higher frequency is more robust to distractors, thus giving them an advantage to entrain the network. Finally, we show that pulsatile inputs can switch between attended inputs in selective attention.

**Author summary:** Oscillations are ubiquitous in the brain and are involved in several cognitive tasks but their role is not completely understood. The Communication Through Coherence theory proposes that background oscillations in the brain regulate the information flow between neural populations. The oscillators that are properly phase-locked so that inputs arrive at the peaks of excitability of the receiving population communicate effectively. In this paper, we study the emerging phase-locking patterns of a network generating PING oscillations under external periodic forcing, simulating the oscillatory input from other neural groups. We identify the conditions for optimal phase-locking and selective communication. Namely, we find that inputs with higher frequency and coherence have an adavantage to entrain the network and we quantify how robust are to distractors. Furthermore, we show how selective attention can be implemented by means of phase locking and we show that pulsatile inputs can switch between attended inputs.

## 1 Introduction

Macroscopic oscillations are widely observed in the brain and they span a temporal scale that ranges from a few to a hundred hertz [1]. There are several studies that associate these rhythms to different cognitive tasks, but its physiological origin and functional role is not completely understood and constitutes an active area of research [2, 3, 4, 5].

Amongst these rhythms, oscillations in the gamma frequency band (30-100 Hz) have been identified in many cortical areas and across different species in a variety of tasks, including attention and memory [5, 6]. Investigations on the mechanisms generating gamma-band oscillations have identified a key role for interneurons [4, 7], which are responsible of generating synaptic inhibitory activity that periodically modulates the excitability of the neurons. These rhythmic changes in the neuronal excitability have been hypothesized to regulate the information flow between distant brain areas in a flexible way. Thus, communication through coherence (CTC) theory [8, 9] proposes that an effective transmission of information between two oscillating neuronal groups occurs when the pre-synaptic input of the sending population reaches systematically the post-synaptic (receiving) population at its maximum phase of excitability (when inhibition has cleared out or is at its lowest value). The oscillators must be therefore properly phase-locked to accomplish effective communication. Synchrony provides a dynamic mechanism to modulate the information flow without modifying anatomical connections, resulting in functional connectivity [10, 11]. Out of this theory, it emerges a solid hypothesis to explain how selective communication emerges in the brain, that is the ability of a receiving neuronal group to respond selectively to different input streams [9].

A growing number of experimental studies have tested the CTC predictions [9]. In particular, some studies have linked the phase of the inhibitory population with the modulation of the input gain of the target population [12]. Also, effective connectivity has been linked to the phase relation between gamma rhythms at the pre and postsynaptic neuronal groups [13]. On the other hand, different studies support that visual and motor selective attention can be implemented through the control of the phase and synchronization between populations [14, 15, 16].

There are also several modeling studies approaching different aspects of the CTC hypothesis. These studies are based either on computational simulations of spiking networks [17, 18, 19, 20, 21] or models that admit an analytically more tractable approach such as single neuron models [22, 23], or firing rate models [10, 24]. Other studies compare different approaches such as in [10, 25]. For instance, in [10], spiking and mean firing rate models were considered, but the latter are not an exact derivation of the former. Thus, in order to perform mathematical analysis one needs to assume important simplifications in the models.

In this study, we take advantage of recently developed neural mean field models to test several hypothesis of CTC theory. Indeed, the development of a new generation of neural mass models in the last years have opened the possibility to test CTC hypothesis in realistic models while simplifying the mathematical analysis. These models provide an exact description (in the limit of large networks) of the macroscopic quantities of a spiking network [26, 27, 28]. Here, we consider a spiking network of excitatory and inhibitory cells (E-I network), whose macroscopic activity in terms of mean firing rate and membrane potential can be described exactly by means of a lowdimensional mean-field model [29, 26]. The low-dimensional mean field model permits mathematical analysis using tools from dynamical systems, and thereby prediction and insight into the relevant parameters that characterize the firing properties of the network. These predictions can then be tested in simulations of the full spiking network. This approach allows us to go beyond our results on CTC obtained in an earlier study for Wilson-Cowan models [24] by considering more realistic networks.

Specifically, we consider a mutually interconnected network of Excitatory and Inhibitory neurons (E-I network), accounting for macroscopic oscillations via the classical PING (Pyramidal Interneuronal Network Gamma) mechanism [4, 7, 30]. These oscillations describe rhythmic changes in the excitability of the E-cells. We apply to this network an external oscillatory periodic input, modeling the input from the pre-synaptic population, for which we modulate the amplitude, the frequency and coherence (that is, how concentrated input volleys are to particular phases of the oscillation cycle).

The present study requires first to examine how a neural oscillator responds to incoming perturbations and the mathematical tool to characterize it is the *phase response curve* (PRC) [31, 32]. The weak coupling assumption enables us to reduce the dynamics close to the oscillator to a single equation, known as the *phase equation* [33]. Since the perturbation is periodic, we define the so-called *stroboscopic map* for the phase equation, whose fixed and periodic points correspond to different phase-locking states between the input and the target network. A bifurcation analysis of the stroboscopic map determines the Arnold tongues, that is, the regions in the parameter space of amplitude and frequency of the periodic forcing, where the different phaselocked states are located.

We explore in detail the solutions that emerge in the phase-locking regions to quantify the communication efficiency in the CTC context. We find that the presynaptic population can entrain the network at different frequencies, and that the range of frequencies is larger for high coherent inputs. Interestingly, we observe that inputs with higher frequency than the natural cycle one establish a stronger effective communication. Moreover, we explore the effects of a disruptor onto the phaselocking and we observe that those inputs with higher frequency are more robust to disturbances, irrespective of its coherence. As a result, the frequency relationship is crucial to prevent desynchronization by disruptors. Finally, we give particular attention to show that a pulse delivered at the appropriate phase can drive selective attention between two similar oscillatory input streams.

## 2 Results

### 2.1 Mathematical Setting for CTC

In this Section we describe the mathematical model for a canonical cortical neural network with intrinsic oscillatory activity in the gamma range. This model will be used to study the effects of an oscillatory input (modeling the input from an emitting population) onto the network activity. In particular, we will explore phaselocking properties between emitting and receiving populations, and its implications for Communication Through Coherence (CTC) theory.

Long-lasting studies on gamma-frequency oscillations have provided the fundamental neurological requirements to generate them: either from the interplay of excitatory (E) and inhibitory (I) populations or from a single inhibitory population with self-feedback. These two processes are referred to as PING (Pyramidal Interneuron Network Gamma) and ING (Interneuron Network Gamma) mechanisms, respectively, and represent the classical models to generate and explain gamma oscillations [7, 4]. In both mechanisms, inhibition plays a strong role. In this paper we focus on the PING mechanism.

To implement the PING mechanism we consider two populations of neurons, one of excitatory (pyramidal) cells and the other one of inhibitory (interneuron) cells, interconnected synaptically. With sufficiently strong and persistent external excitatory drive to the pyramidal neurons, the E-cells activate and recruit the Icells which, in turn, send a reciprocal inhibitory feedback onto the E-cells. The inhibition causes the E-population to be less receptive to the external drive until its effect decays, starting again the E-I cycle. This mechanism generates a self-sustained macroscopic oscillatory activity in the gamma range (see Figure 1).

**Figure 1:**
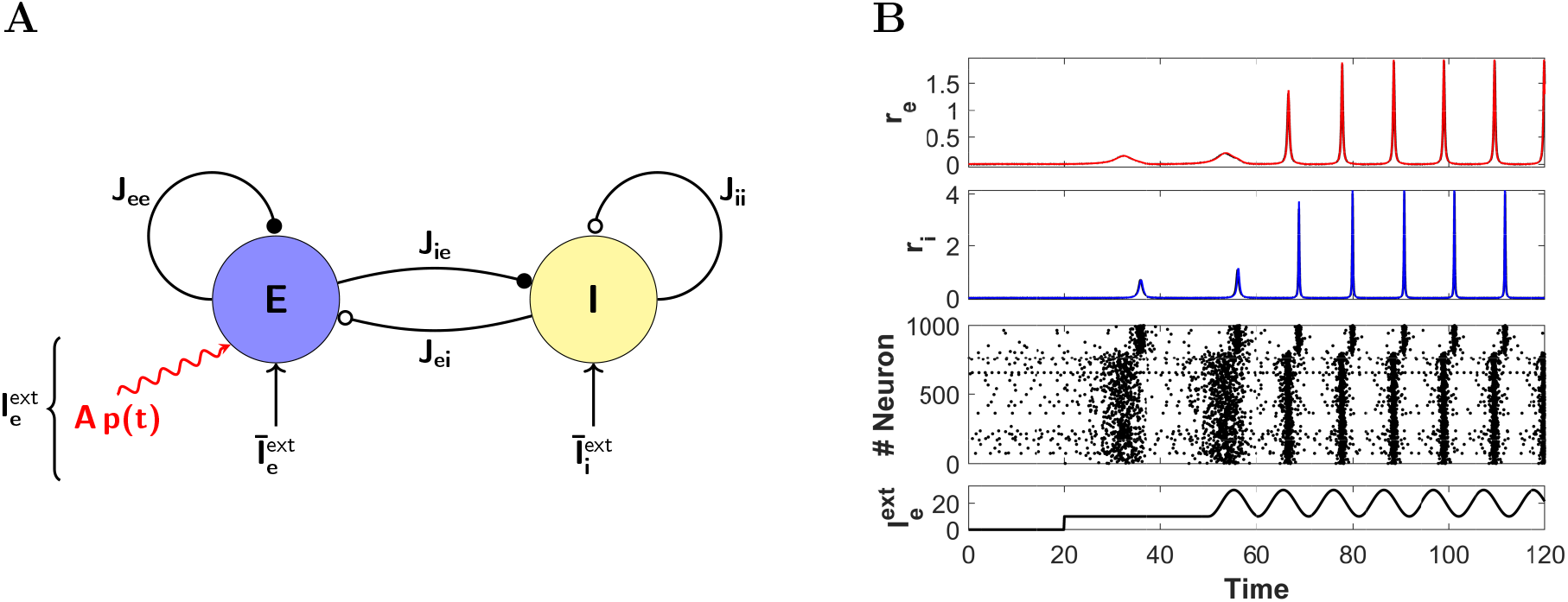
E-I cortical network and comparison between full network and reduced system. (A) Schematic representation of a cortical neural network consisting of excitatory (E) and inhibitory (I) cells. Excitatory (resp. inhibitory) synapses are depicted by arrows with a black-filled (resp. empty) circle pointing at the receiving population. The parameter J_ab_, *a*, *b* ∈ {*e, i*} is the connectivity strength of the *b* → *a* synapse. Besides synaptic input, each population also receives an external tonic current 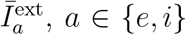. Additionally, the E-population receives a periodic input *Ap(t)* (red curve), modeling the input from an oscillating neuronal population. (B) From bottom to top: Time evolution of the external input current 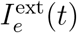 onto the E-cells. Raster plot of 1000 randomly selected neurons (the first 800 neurons are excitatory and the last 200 inhibitory). Time evolution of the macroscopic quantities *r_e_* and *r_i_* obtained from simulations of the mean-field model (1)-(2) (red and blue curves, respectively) and the averaged firing rate activity of the full spiking network (black). In the latter case, the mean firing rate has been computed by averaging the number of spikes in a time window of size *δt* = 8 · 10^−2^ Parameters : *N_e_* = *N_i_* = 5000, 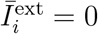 and the rest of the parameters as in (6)

To model the E-I network we consider a neural mass model that was recently derived for networks of spiking neurons [26, 29]. It provides an exact description (in the thermodynamic limit) of the macroscopic quantities of a spiking network of quadratic integrate-and-fire neurons, namely, the mean firing rate *r* and the mean membrane potential *V* (see Methods for more details).

The model is given by a set of differential equations for the E-population,

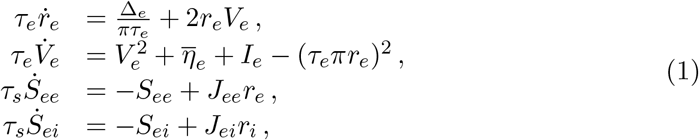

and another identical set for the I-population,

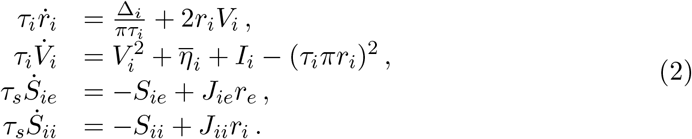

Here, *V_a_* and *r_a_* represent the mean voltage and mean firing rate of the population *a* ∈ {*e*, *i*}. The parameters *η_a_* and *Δ_a_* are, respectively, the center and width of the static distribution of inputs to the individual spiking neurons, which is considered to be Lorentzian. The time constant *τ_a_* is the membrane time constant of the individual neurons in population *a*. The variable *S_ab_* models the synaptic current from the presynaptic population *b* to the postsynaptic population *a*, and *J_ab_* is the strength of the corresponding synaptic connection. The time constant **τ_s_** is the synaptic time constant.

The total input current to the excitatory population is given by

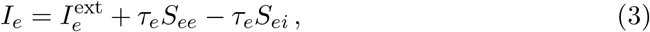

and to the inhibitory one, by

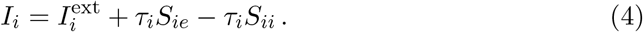

Notice that the terms *I_e_* and *I_i_* above provide the coupling between systems (1) and (2). The terms 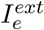 and 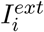 represent the external input drive.

In this study, we are going to consider that the input current to the I-cells 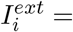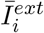 is a tonic current, while for the E-cells might be time dependent, that is,

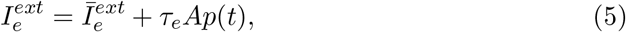

where 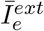 is a tonic current and *p*(*t*) is a time periodic function. Here, A is a parameter that modulates the amplitude of periodic input *p*(*t*). We are going to consider different functions for *p*(*t*) (see Methods) to explore different phase-locking properties. Notice the time constant *τ_e_* multiplying the periodic input.

For this paper, we consider fixed the following parameters for system (1)-(2) (they are the same values as for the PING setting in [29]):

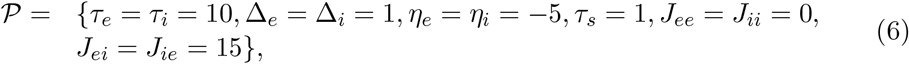

and we will vary the other ones.

In Figure 1A we present a schematic representation of the E-I neural network described by system (1)-(2) and their excitatory and inhibitory synaptic connections. In Figure 1B we show numerical simulations of the firing rate activity of the network in response to a time dependent input 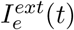 using the mean field reduced model (1)-(2) and the full spiking network model. Notice the perfect agreement between both models. We emphasize here the power of this reduced description, which will allow us to perform the mathematical analysis detailed in the next Sections.

#### 2.1.1 Characterization of gamma oscillations

Before studying the effects of a periodic perturbation onto the E-I network, we characterize the network intrinsic oscillatory activity, that is, when *A* = 0 in (5) and 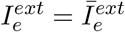 is a tonic current.

We first identify the values of the external inputs 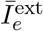 and 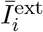 for which the model (1)-(2) shows oscillations. With strong enough constant drive to the E-cells, the system (1)-(2) transitions from an asynchronous state of low activity (resting) to an oscillatory regime (see for instance the transition at *t* = 20 in Figure 1B). From the viewpoint of dynamical systems, we say that the system undergoes a bifurcation – a qualitative change in the underlying dynamics when a parameter is varied (in our case when 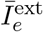 or 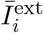 are varied).

We identify two relevant bifurcations in the 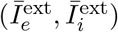-plane to determine the oscillatory states of the system (see Figure 2A). The asynchronous activity states destabilize through Hopf bifurcations (blue curve), which give rise to oscillations (periodic orbits) that can be stable or unstable depending on whether the Hopf bifurcation is supercritical (solid blue curve) or subcritical (dashed blue curve). Along this curve, Generalized Hopf (GH) bifurcations (also known as Bautin bifurcations) occur at the places (red circles) where the Hopf bifurcation changes from supercritical to subcritical (or vice versa) [34]. Moreover, oscillations also disappear (resp. appear) through saddle node (or fold) bifurcations of periodic orbits (purple curve). At these bifurcations, a stable and an unstable periodic orbit collide and annihilate each other (resp. sudden create). Thus, oscillations occur in the blue and orange regions enclosed by the Hopf and saddle node bifurcations. Remarkably, in the orange region, the system presents bistability between asynchronous activity state and oscillatory state. There is also a small region (intersection of orange and blue regions) showing bistability between two different types of oscillations. Studying the dynamics in the bistable regions is a topic of interest for understanding the role of oscillations in cognitive tasks. See for instance [35] for a study of the response of multistable networks of recurrently coupled spiking neurons to external oscillatory drive and its implications for working memory and memory recall tasks. However, the topic is beyond the scope of this paper and is left for future research.

**Figure 2:**
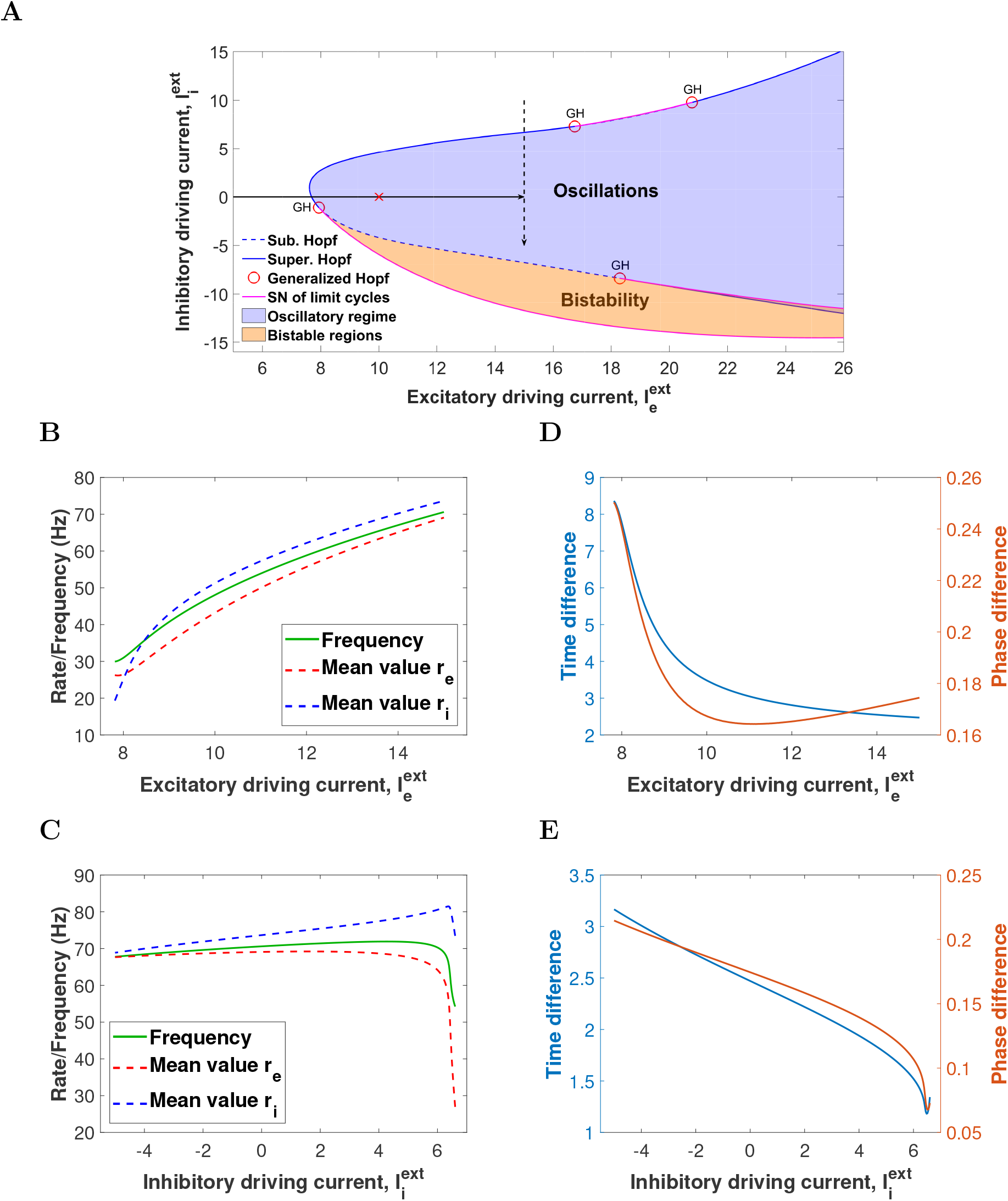
E-I cortical network with oscillatory activity in the gamma range. (A) Two-parameter bifurcation diagram of system (1)-(2) for the excitatory current 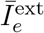 (x-axis) and the inhibitory current 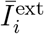 (y-axis). The solid (resp. dashed) blue curve corresponds to a supercritical (resp. subcritical) Hopf bifurcation curve. Red circles correspond to (codimension 2) Generalized Hopf bifurcations. Purple curve corresponds to saddle node bifurcation of limit cycles. Oscillations occur in the blue and orange regions. Red cross corresponds to values 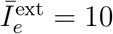 and 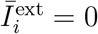 generating the limit cycle considered later on. (B, C) Frequency oscillation (green) and integral mean values of the firing rates *r_e_* (dashed red) and *r_i_* (dashed blue) as a function of (B) tonic excitatory current 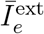 and (C) tonic inhibitory current 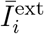. See equation (7). (D, E) Time difference (blue) and relative phase (orange) between inhibition and excitation as a function of (D) tonic excitatory current 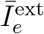 and (E) tonic inhibitory current 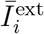. In both panels B and D the inhibitory current 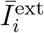 is set to 0. In Panels C and E the tonic excitatory current 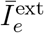 is set to 15. Other parameters are as in (6).

To characterize the oscillations, we compute their frequency and the integral mean values of the firing rate of both populations, that is,

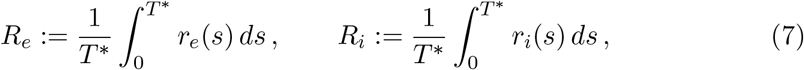

where *T* ^*^ is the oscillation period. The values *R_e_* and *R_i_* in (7) provide an average of the firing rate of the individual E and I-cells, respectively.

Moreover, we compute the time difference between the maximum of the I and E firing rates in one cycle (I to E latency) and the relative phase, i.e., the time difference normalized by the period. These factors provide information about the time it takes for the I-cells to activate and halt the activity of the E-cells, and suggest possible gates for communication in the CTC context [25].

We first consider a fixed value of the external inhibitory current 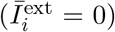, and we increase the excitatory one 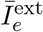 from 0 to 15. This interval covers the progression towards the oscillatory region (see solid arrow in Figure 2A). In Figure 2B we show the frequency of the emerging oscillations as a function of the current injected into the E-cells (green curve). Notice that the frequency is bounded away from zero (characteristic of a Hopf bifurcation) and increases from 30 to 70Hz, which fits in the gamma range. The average quantities (7) are close to the frequency oscillation curve for all values of the external current 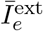 that we have explored. Thus, the average firing rate of the E-I cells is close to the macroscopic oscillation frequency, and slightly higher for the I-cells than for the E-cells. Increasing the external input onto the E-cells causes only a slight shortening of the latency period of inhibition over excitation, especially for oscillations above 40Hz (see Figure 2D).

We also vary the constant external drive onto the inhibitory cells 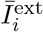 from 10 to −5 (notice that it may take negative values), while the input to the E cells is kept fixed at 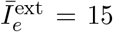. The system is initially in the non-oscillatory region and enters the oscillatory region (see dashed arrow in Figure 2A). As before, the integral mean firing rates *R_e_* and *R_i_* (dashed lines in Figure 2C) remain close to the frequency of the macroscopic oscillations (green line in Figure 2C). Increasing the depolarizing current onto the I-cells 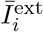 does not significantly affect the macroscopic oscillation frequency (the frequency remains close to 70 Hz, except for the final drop around 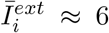, in contrast with the effect of a depolarizing current onto the E-cells. However, increasing 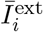 causes a strong effect on the inhibition-to-excitation latency (compare phase difference with Figure 2D), the inactive period after inhibition acts on excitation shortens, and thus the relative phase (see Figure 2E).

In summary, the oscillation frequency is governed by a trade off between the decay time of the inhibitory conductance and the external excitatory drive onto the E-cells. Thus, increasing the depolarizing drive onto the E-cells 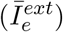 shortens the oscillation period, because excitation can overcome the effect of the inhibition sooner. On the other hand, increasing the depolarizing drive onto the I-cells 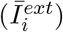, activates the I population faster, thus shortening the E-I latency.

Next, we are going to study the different phase-locking patterns that emerge when the oscillatory neural E-I network receives a periodic external input. To do so, we consider the oscillations that emerge for values 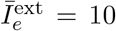 and 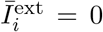 (red cross in Figure 2A). Notice that for these values, individual E-cells spike, on average, less frequently than I-cells (compare *R_e_* and *R_i_* in Figure 2B for 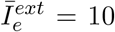). However, they both have firing rates similar to the oscillation frequency of the network, which is about 48Hz (period *T* ^*^ = 20.811 ms), clearly in the gamma range. Moreover, the time difference between the E and I volley peaks is short (around 4ms, see Figure 2D). Figure 3A shows the evolution of the variables of the system along this periodic orbit. Later in Section 2.6, we will explore another type of oscillations, closer to the Hopf bifurcation curve, showing different features.

**Figure 3:**
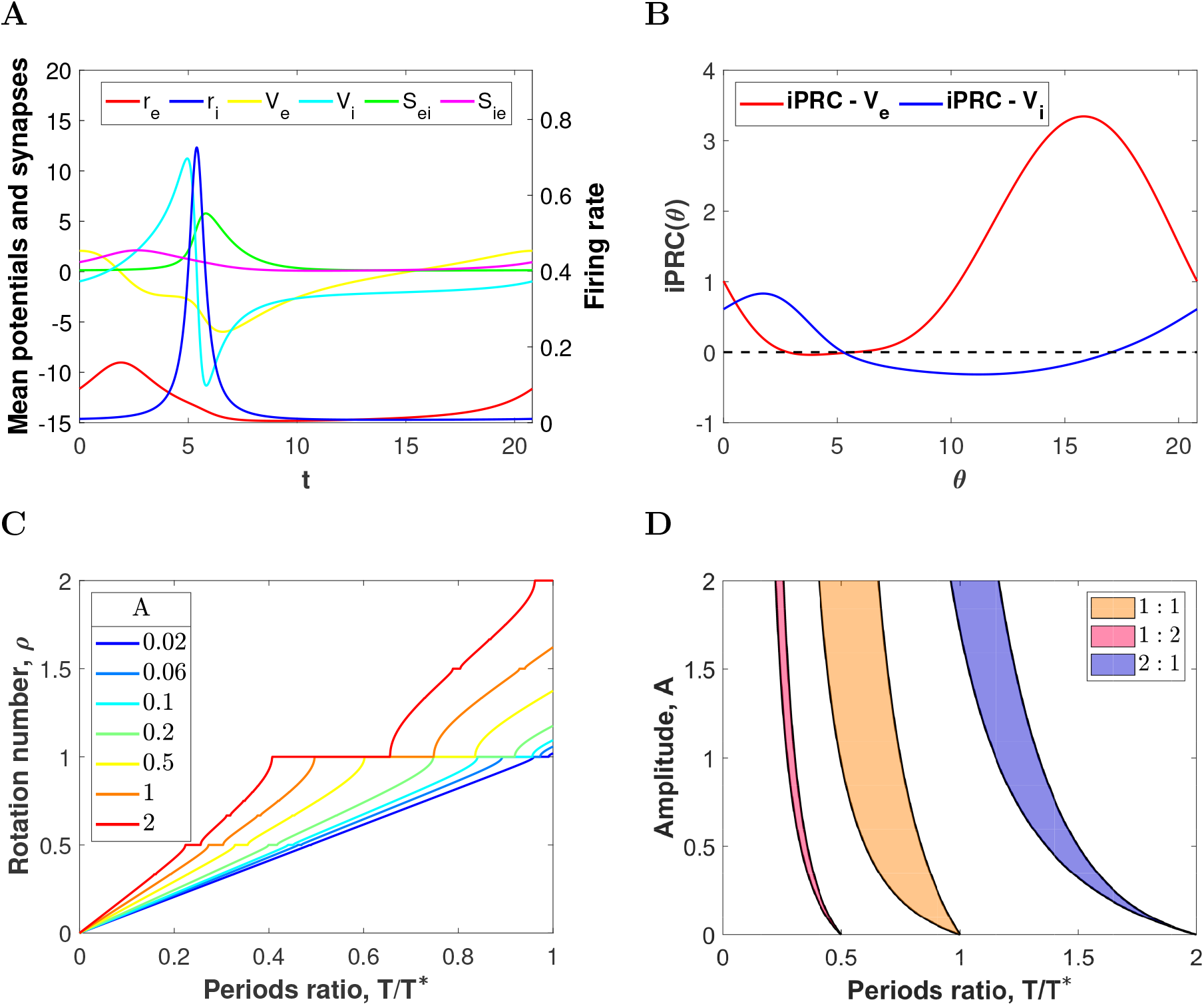
Rotation numbers and phase-locking regions for the sinusoidally forced PING oscillation. (A) Temporal evolution of firing rate, mean membrane potential and synaptic variables over a cycle of a PING oscillation corresponding to the red cross in Figure 2A. (B) Infinitesimal Phase Response Curve (iPRC) for perturbations in the direction of the variables *V_e_* and *V_i_* (red and blue curves, respectively). (C) Rotation numbers of the stroboscopic map (9) for a sinusoidal perturbation (15) applied in the direction of *V_e_*, as a function of the ratio between the intrinsic period of the E-I network *T* ^*^ and the input period *T* for different values of the amplitude *A*. (D) Arnold tongues (up to *A* = 2) corresponding to the most prominent phase-locked states in Panel C: 1:1 phase-locking (orange), 1:2 phase-locking (purple) and 2:1 phase-locking (blue).

### 2.2 Mathematical Analysis of Phase Dynamics

In this section we describe the mathematical analysis used to study the phase relationship between the neural oscillator described by the low dimensional firing-rate model (1)-(2) and an external time periodic stimulus *p*(*t*) (see equation (5)).

#### 2.2.1 The phase equation

Consider the case for which the E-I network is showing macroscopic oscillatory activity by means of a PING mechanism. Then, for an adequate set of parameters (see previous Section), the system of differential equations (1)-(2) has a hyperbolic asymptotically stable limit cycle Γ, of period *T* ^*^, attracting all nearby orbits. This stable limit cycle is referred to as the oscillator.

Any oscillator can be parametrized by a single phase variable *θ* measuring the elapsed time (modulo *T* ^*^) from a reference point on the limit cycle, pinpointed as *θ*_0_ (i.e. *θ*(*t*) = *t* + *θ*_0_ mod *T* ^*^). From now on, we will assume *θ*_0_ = 0. The phase variable *θ* describes uniquely a point on the oscillator via the periodic solution, i.e. *γ*(*θ*(*t*)) = *γ*(*t*) = (*r_e_*(*t*), *V_e_*(*t*), … , *r_i_*(*t*), *V_i_*(*t*), …), such that *γ*(*t*) = *γ*(*t* + *T* ^*^).

When we apply an external periodic input *p*(*t*) of period *T* (see equation (5)) through the dynamics of the variable *V_e_*, the evolution of the phase variable *θ* is described by the equation,

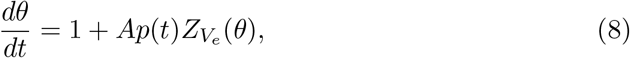

where 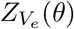 is the infinitesimal Phase Response Curve (iPRC) in the direction of the variable *V_e_*. The iPRC measures the oscillator’s phase shift due to an infinitesimal perturbation applied to the voltage *V_e_* at different phases of the cycle [31]. The positive parameter A controls the strength of the input, and it is assumed to be sufficiently small (A « 1). See Methods. In Figure 3B, we show the iPRC 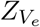 (red curve) for the oscillator in Figure 3A (corresponding to the parameter set 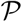 in (6) with 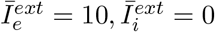). Notice that the iPRC 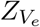 is almost input insensitive (close to zero) for phases of the cycle corresponding to the activation of the I-volley, and positive for the rest, achieving its maximum value right before the activation of the E-volley. The fact that the iPRC is mainly positive indicates that this oscillator can only advance the phase, and therefore synchronize with higher frequency inputs.

We also include the iPRC when the infinitesimal perturbation acts on the voltage *V_i_* (blue curve). Later in Section 2.6, we will explore another type of oscillations, closer to the Hopf bifurcation curve, for which the iPRC takes both positive and negative values.

#### 2.2.2 Stroboscopic map and phase-locking

Since the input *p*(*t*) is periodic of period *T*, we use the so-called *stroboscopic map* to study the phase equation (8). Let *Φ_t_*(*θ*) be the solution of the differential equation (8) starting at phase *θ*_0_ at *t* = 0. The stroboscopic map *P* (also known as Poincaré phase map) is defined on the circle and provides the phase of the oscillator after a time *T* (the period of the perturbation), that is

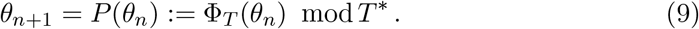

The dynamics of the stroboscopic map provides information on phase-locking between the oscillator and the periodic perturbation. In our context, we say that they are *p* : *q phase-locked* if every time the forced oscillator completes *p* revolutions, the oscillatory input completes *q* revolutions. The concept of synchronization is often associated to the 1:1 phase-locking, that is, the external stimulus entrains the oscillator and both oscillate at the frequency of the forcing.

The 1:1 phase-locked states correspond to fixed points of the stroboscopic map *P* in (9), because, after integrating the *phase equation* for a time *T* (the period of the forcing), the phase of the forced oscillator coincides with the initial one (modulo *T* ^*^), i.e. *θ* + *T* ^*^ = *P* (*θ*). The relation *p* : 1, consisting in *p* revolutions of the oscillator per one of the stimulus, also corresponds to a fixed point of the stroboscopic map (we must find a solution of *θ* + *pT* ^*^ = *P* (*θ*)). Finally, in the most general framework, *p* : *q phase-locked* states correspond to *q*-periodic points of the stroboscopic map, i.e. solutions of the equation *θ* + *pT* ^*^ = *P* ^*q*^(*θ*).

#### 2.2.3 Rotation number and Arnold tongues

To identify the fixed and periodic points of the stroboscopic map *P* in (9) (and therefore, the phase-locked states) we compute the rotation number. The rotation number (denoted by *ρ*) measures the angle on the circle that the map *P* turns on average at each iterate. See Methods for the formal definition. Results in dynamical systems theory [36] guarantee that whenever *ρ* is rational (i.e. *ρ* = *p/q* for *p*, *q* ∈ ℕ), there exists a *q*-periodic point of the stroboscopic map (9), that is, a solution of *P* ^*q*^(*θ*) = *θ* mod *T* ^*^, corresponding to a *p* : *q* phase-locking relationship. If, by contrast, *ρ* is irrational, then the orbits of (9) fill densely the circle, and phase-locking does not occur.

We have computed the rotation number for the stroboscopic map (9) of the phase equation (8) corresponding to the oscillator in Figure 3A (whose iPRC 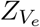 is shown in Figure 3B) and a depolarizing input *p*(*t*) of sinusoidal type (see equation (15)), for which we vary the frequency (1/*T*) and the amplitude, controlled by the parameter *A*.

When *A* = 0, the only *p* : *q phase-locked* states occur at rational values of the frequency relationship, *T/T* ^*^ = *p/q*. However, as *A* increases there appear plateaus in the graph of the rotation number as a function of *T/T* ^*^, which is known as *Devil’s staircase*. The plateaus, corresponding to rational values of the rotation number, appear because there is a range of periods *T* close to 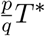 that also give rise to *p* : *q* phase-lockings (see Figure 3C). The easiest detectable plateaus correspond to *ρ* = 1, 1/2, 2 for the range of input frequencies considered. Indeed, for small amplitudes *A* = 0.02 (dark blue curve), we observe one small plateau at height 1 for periods *T* similar to the natural one *T* ^*^ (i.e. *T/T* ^*^ ≈ 0.95), corresponding to 1:1 phase-locking. Similarly, at amplitude *A* = 0.06 a (very) small plateau at height 1/2 becomes visible around *T/T* ^*^ ≈ 0.45, corresponding to 1:2 phase-locking. Increasing the amplitude, these plateus are lengthened and shifted towards higher input frequencies. It is also remarkable the emergence of other relevant phase-locked states such as the 2:3, 3:2 and the 2:1 (see plateaus at height 2/3, 3/2 and 2, respectively, for *A* = 2).

In the relative frequency vs amplitude parameter space (*T/T* ^*^, *A*)-plane, a *p* : *q* phase-locking region resembles a leaf or a fang with their tip stuck at *A* = 0, known as *Arnold tongue* [36] (see Figure 3D). The plateaus in the graph of the rotation number are actually slices of such Arnold tongues obtained by fixing the amplitude (see Figure 3C). The shape of these Arnold tongues is strongly linked to the sign of the iPRCs. Indeed, when the iPRC is mainly positive (e.g. the iPRC 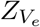 in Figure 3B), the oscillator can essentially only advance its phase and thus it synchronizes almost exclusively to fast external oscillations, that is, periodic forcing with period *T* < *T* ^*^. The resulting Arnold tongue is bent to the left (see Figure 3D). This fact justifies why the *p/q* plateaus in Figure 3C were shifted gradually to the left (see, for instance, at the 1:1 phase-locked state for different amplitudes).

### 2.3 Inputs with higher frequency communicate more effectively (sinusoidal case)

Phase-locked states have implications for neuronal communication in the CTC context. Indeed, the CTC theory proposes that neural communication between two populations is more effective when the underlying oscillations are properly phase-locked so that windows for sending and receiving are open at the same time [8, 9]. In the unidirectional setting considered herein (see Figure 1A), if the external input volley reaches the receiving E-population before the inhibition activates, it may contribute to increase the activity of the E-cells, thus promoting effective communication. On the other hand, if the input volley arrives to the E-population when the inhibition is present, then such a stimulus may not have any effect on the activity of the E-cells. Hence, effective communication depends essentially on the timing of the inhibition and the external input [37].

In the previous section, we have identified several phase-locked states between an oscillating E-I network (PING interplay) and a sinusoidal input using the phase reduction (see Figure 3). To characterize quantitatively whether the phase-locking that emerges is optimal for communication, we compute the factor Δ*τ* (equation (24)), which measures the time difference between the peaks of the input and the I-volley, and the factors 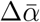, Δ*α* and Δ*γ* which measure the firing rate response of the E-cells of target network to the input (see Methods). More precisely, 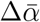 (equation (25)) provides information about the rate change in the overall firing rate of the E-population over the whole cycle, Δ*α* (equation (26)) about the rate change in the maximum of the E-volley and Δ*γ* (equation (27)) about the rate change in the half-width of the E-volley. In our previous work [24], we considered only the factor Δ*α*, but the quantities 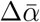 and Δ*γ* allow us to complement the information about the changes in the firing rate. In particular, we can explore whether “communication” means that the target network shows higher firing rates and/or higher coherence, that is, enhancement of synchronization in spiking neurons within the target population [21].

We have computed these factors for the periodic solutions of system (1)-(2) perturbed with a sinusoidal input *p*(*t*) defined in (15), in the 1:1, 1:2 and 2:1 phaselocking regions (identified in Figure 3), where the existence of these orbits is guaranteed. Recall though that the computation of the phase-locking regions in Figure 3 was carried out under the weak coupling hypothesis (small amplitude *A*). For this reason, the prediction of phase-locking might be less accurate as the amplitude increases. Remarkably, computations only show a slight disagreement between the prediction obtained from the phase equation and the solution found for the perturbed 8-dimensional system (1)-(2) (compare the black curve and the colored region in Supplementary Figure S1). This outcome guarantees the reliability of the predictions based on the phase reduction method at least for amplitudes with *A* < 1. In Figure 4 we show the values of the factors Δ*τ*, Δ*α*, 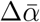 and Δ*γ* along constant amplitude values that range between *A* = 0.1 and *A* = 1 (identified by the color of the curve) in the corresponding Arnold tongue.

**Figure 4:**
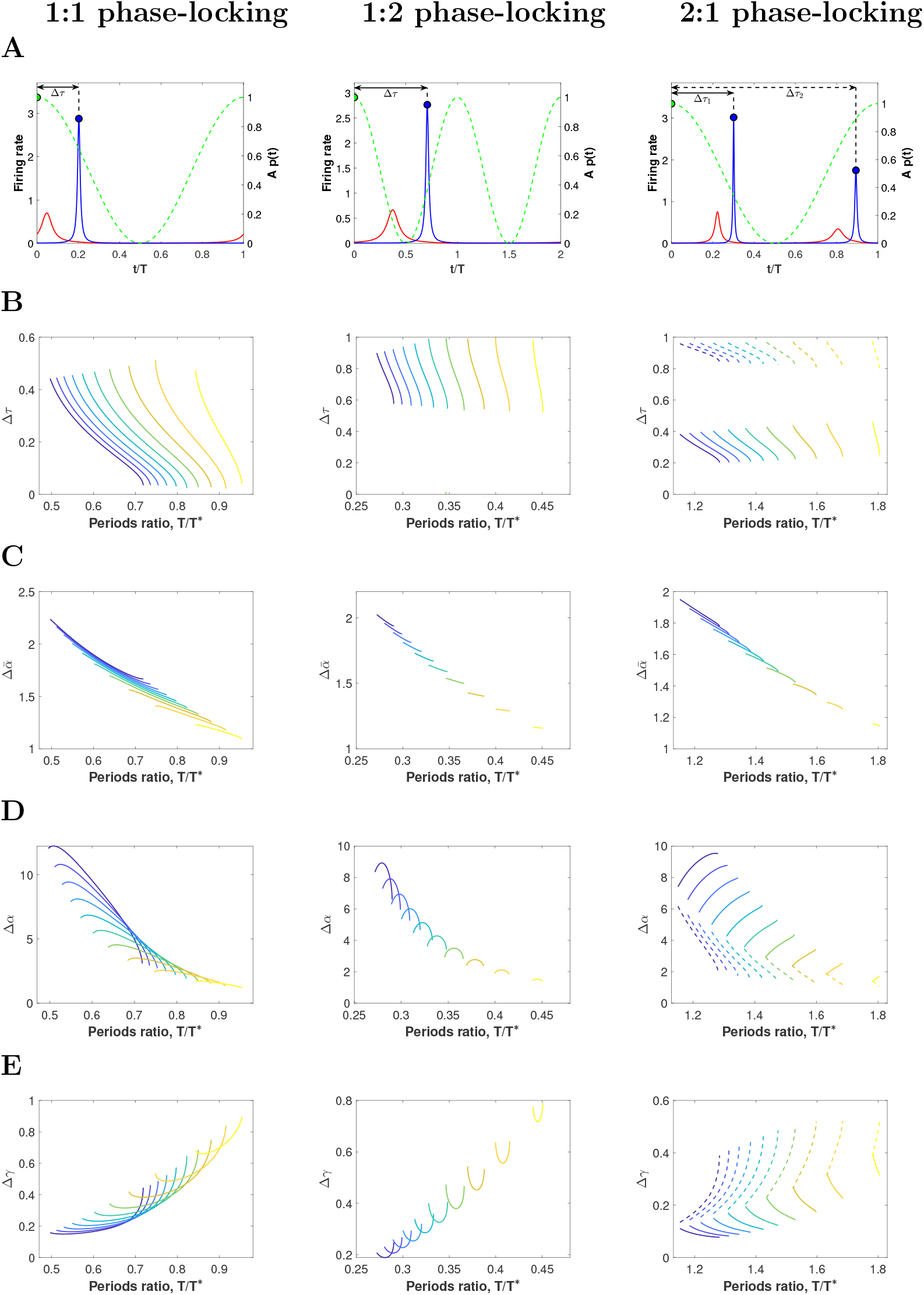
Input effectiveness for network entrained by sinusoidal inputs. (A) Evolution of the firing rate variables *r_e_* (red) and *r_i_* (blue) and the sinusoidal perturbation (dashed green) for a representative periodic orbit within the 1:1 (left), 1:2 (middle) and 2:1 (right) phase-locking regions. (B-E) Factors describing effective communication within these phase-locking regions for orbits calculated along (equidistant) sections *A* = ct of the corresponding Arnold tongues, indicated by the color of the curve (ranging from dark blue, *A* = 1, to yellow, *A* = 0.1, with increments of size 0.1). The factors are: (B) Δ*τ*, describing the timing between inhibition and input volleys (normalized by the input period *T*), (C) 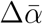, describing the rate change in the averaged firing rate, (D) Δ*α*, describing the rate change in the maximum of the (excitatory) firing rate, and (E) Δ*γ*, describing the rate change in E-volley half-width. See Methods. Additionally, the factors Δ*τ*, Δ*α* and Δ*γ* within the 2:1 Arnold tongue are also shown for the oscillator’s second peak (dashed lines).

We observe that in the 1:1 phase-locking regions (see Figure 4 left), the factor Δ*τ* (see Figure 4A left) indicates that inhibition follows the perturbation (0 < Δ*τ* < 0.5 mainly within the 1:1 Arnold tongue) but Δ*τ* changes dramatically within the Arnold tongue (see Figure 4B left). Indeed, the factor Δ*τ* reaches its maximum on the left boundary of the Arnold tongue and it decreases gradually as the input frequency decreases and approaches the right boundary (*T/T* ^*^ increases). Thus, faster inputs (i.e. *T/T* ^*^ close to the left boundary) arrive as much apart as possible from the inhibition (Δ*τ* ≈ 0.5), while those with frequencies more similar to the intrinsic oscillatory frequency (*T/T* ^*^ close to the right boundary) arrive almost at the same time as the inhibition (Δ*τ* ≈ 0). See also Supplementary Figure S2. Therefore, we observe a greater increase in the activity of the E-cells due to the input (that is, a large 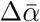) on the left hand side than on the right hand side of the Arnold tongue (see Figure 4C left). Recall that when 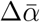 is close to 1 (as it happens close to the right boundary), the effect of the input is almost negligible, so we consider that the input is ignored and effective communication is not established. Remarkably, if the amplitude of the input is large enough, it can overcome the presence of inhibition and elicit a response in the E-cells. So, in conclusion, the firing rate of the E-cells grows with both the amplitude and the frequency of the input.

We emphasize that an optimal phase relationship causes both higher firing rates and higher coherence to the target network. Indeed, increases in the overall firing rate of the target population (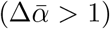), correlate with sharpening the E-volley: the half-width decreases (Δ*γ* < 1) and the maximum increases (Δ*α* > 1). Compare Figures 4C, D and E (left column).

For the 1:2 phase-locking region (see Figure 4 middle), the network receives two input volleys but responds with only one E-I volley every cycle (see, for instance Figure 4A middle). Thus, only one input volley elicits a response in the target network, while the other one is ignored. Here, the factor Δ*τ* (see Figure 4A middle) measures the time distance from the maximum of the I-volley to the preceding input volley and ranges between 0 an 1 (see also Methods). We observe (Figure 4B middle) that inhibition overlaps with one input volley on the left boundary of the Arnold tongue (Δ*τ* ≈ 1) and shifts and locates between both inputs volleys as the frequency decreases (Δ*τ* decreases down to 0.5 on the right boundary of the 1:2 Arnold tongue). See also Supplementary Figure S3. Consequently, the factor 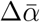, measuring changes in the overall firing rate, is larger on the left boundary than on the right one (see Figure 4C middle). As before, the response of the E-cells also grows with the amplitude of the input. Interestingly, the E-volley is sharper in the center of the Arnold tongue than close to the boundaries (see non-monotonic shape for Δ*α* and Δ*γ* in Figure 4D and E). This non-monotonic shape is also slightly observed at the left boundary of the 1:1 Arnold tongue. At the left boundary, the input frequency is high relative to the inhibition time constant governing the natural period of gamma oscillations, thus inhibition starts to step on the input volley because there is not enough time to wear off its effects. Indeed, when the input frequency is increased a bit more, phase-locking is lost. Later on, we will see that this non-monotonic shape is more pronounced for high coherent inputs.

In the 2:1 phase-locking region (see Figure 4 right), there are two E/I-volleys of the network per one of the input (see Figure 4A right). That is, the input is slower than the intrinsic oscillatory activity of the network. We compute the factor Δ*τ* for each I-volley (see Figure 4A right). One of the I-volleys arrives after the maximum of the input (0 < Δ*τ*_1_ < 0.5, see solid curve in Figure 4B right), while the other one ahead of it (0.5 < Δ*τ*_2_ < 1, see dashed curve in Figure 4B right). See also Supplementary Figure S4. Consequently, the response of the E-cells is enhanced when inhibition follows the input peak (solid curve in Figure 4D and E right) than when it precedes it (dashed curve). Interestingly, on the left boundary of the 2:1 Arnold tongue the increase in the overall firing rate is higher (see 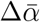 in Figure 4C right) and the external input modulates both E-volleys similarly (solid and dashed curves for Δ*α* and Δ*γ* in Figure 4D and E right almost touch). However, on the right boundary of the 2:1 Arnold tongue, the overall firing rate increase is smaller (see 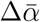 in Figure 4C right), but the differences in the modulation of the maximum and half-width of the E-volleys are more pronounced (see dashed and solid curves separate in Figure 4D an E right). Thus, the low frequency input can be seen as a modulator of the E/I-network rhythm (see Discussion).

### 2.4 Increasing the coherence of the periodic input enlarges the frequency range that entrains the network

In this section, we explore how the results obtained for sinusoidal inputs extend to more realistic inputs. In particular, we consider a periodic input for which we can vary its coherence. In this paper, the concept of coherence refers to concentration of the periodic input volley around some phases of the cycle (as in [22]). To that end, we use a periodic input where each cycle is described by the von Mises probability density function, which has the expression given in (17) (see Methods). Its coherence is modulated by the parameter *κ* in formula (17), while the area over a cycle remains constant independently of *κ*, that is, the total external drive is maintained (see Figure 5A and Methods).

**Figure 5:**
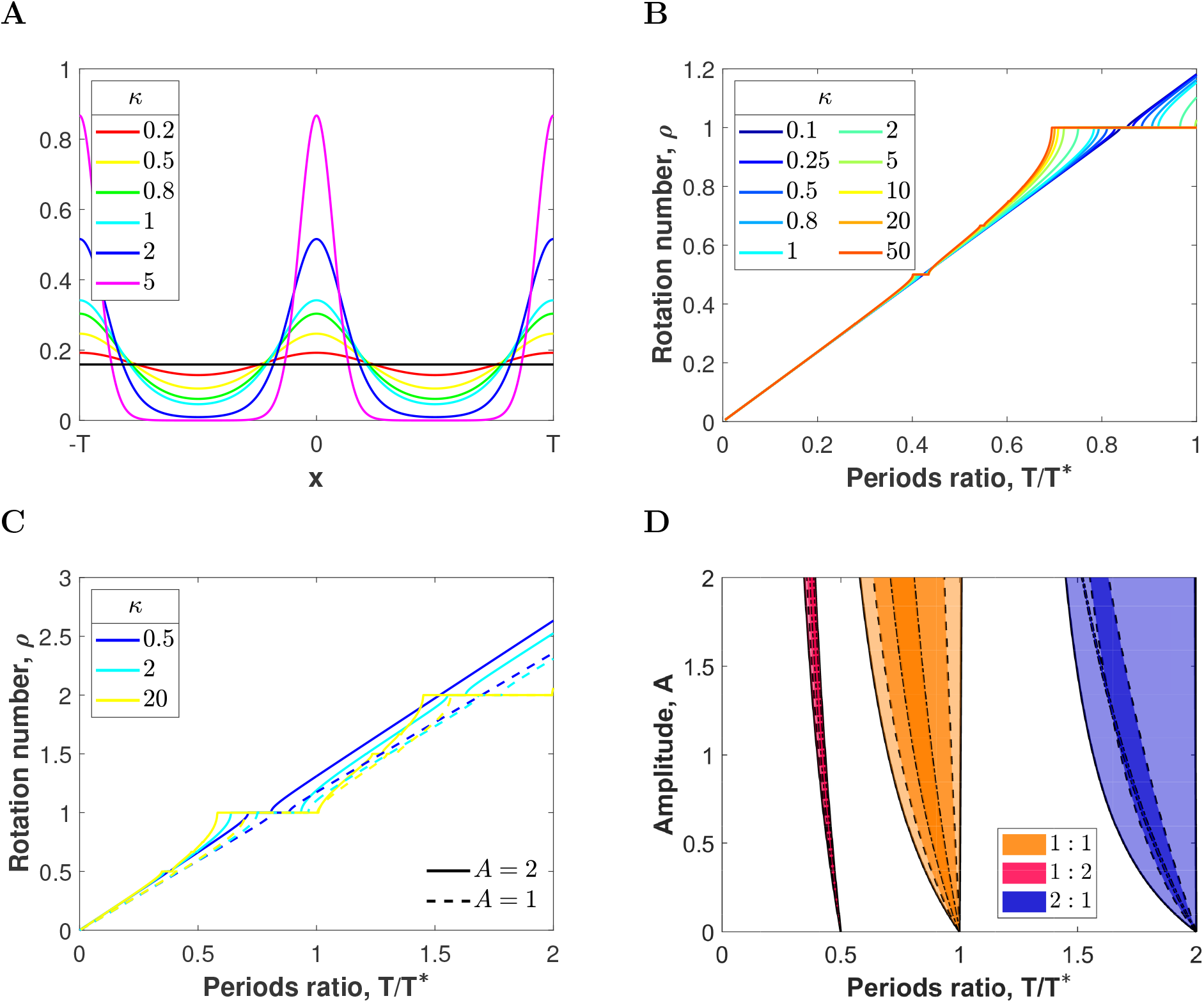
Rotation numbers and phase-locking regions for the forced PING oscillation varying input coherence. (A) Von Mises (Circular) distribution as a function of the factor *κ* controlling the input coherence. Large values of *κ* result in distributions concentrated around the location *μ* = 0, whereas smaller values lead to broader low-amplitude distributions. The black horizontal line corresponds to the uniform distribution (limit case attained when *κ* = 0). (B,C) Rotation numbers of the stroboscopic map (9) for a von Mises input (16) applied in the direction of *V_e_*, as a function of the ratio between the intrinsic period of the E-I network *T* ^*^ and the input period *T*. (B) Rotation numbers for *A* = 1 and different input coherence values *κ*. Broader, low-amplitude inputs (*κ* < 1) lead to small 1:1 and 1:2 plateaus, which lengthen when the external input is more coherent (*κ* 1). (C) Rotation numbers up to *T/T* ^*^ = 2 (comprising both faster and slower inputs) for three representative values of *κ* (low, medium and high coherence) and for amplitudes 1 (dashed lines) and 2 (solid lines). Increasing the amplitude shows a more pronounced shift of the plateaus. (D) Arnold tongues (up to *A* = 2) corresponding to the 1:1 (orange), 1:2 (purple) and 2:1 (blue) phase-locked states for different input coherence: *κ* = 20, 2, 0.5 corresponding to the regions delimited by solid, dashed and dash-dotted curves, respectively.

As for the sinusoidal input, we have computed the rotation number and Arnold tongues following the methodology described in Section 2.2 (see Figure 5). We observe that higher coherent inputs (larger *κ*) give rise to devil’s staircases with larger rational plateaus (compare the red curves with the blue ones in Figure 5B). Moreover, when looking at the Arnold tongues of the largest phase-locking regions in the devil’s staircases (1:1, 1:2 and 2:1), we observe that lower coherence inputs (small *κ*) result in narrower Arnold tongues, enclosed in those with larger *κ* (see Figure 5D). Thus, inputs with higher coherence are capable to entrain the target network for a wider range of frequency values. Notice that, as before, Arnold tongues are bent towards the left on the (*T/T* ^*^, *A*)-parameter space, as it is expected, because the iPRC is mostly positive (see Figure 3B). Noticeably, as input coherence increases, the right boundary of the Arnold tongues is set approximately at a fixed frequency relationship *T/T* ^*^ ≈ *p/q* independently of the amplitude *A* (notice the right boundary of the 1:1 and 2:1 Arnold tongues for *κ* = 20 is fairly vertical, plotted in Figure 5D). Thus, increasing the input coherence enlarges the range of frequencies that entrain the network without shifting it.

Next, we explore whether the type of entrainment that emerges inside the 1:1, 1:2 and 2:1 phase-locking regions for high coherence inputs (*κ* = 20) is optimal for communication in the CTC context. To do so, we compute the factors Δ*τ*, 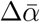, Δ*α*, and Δ*γ*, described in the previous section, for periodic orbits of the perturbed system (1)-(2) within the corresponding Arnold tongue. Results are shown in Figure 6.

**Figure 6:**
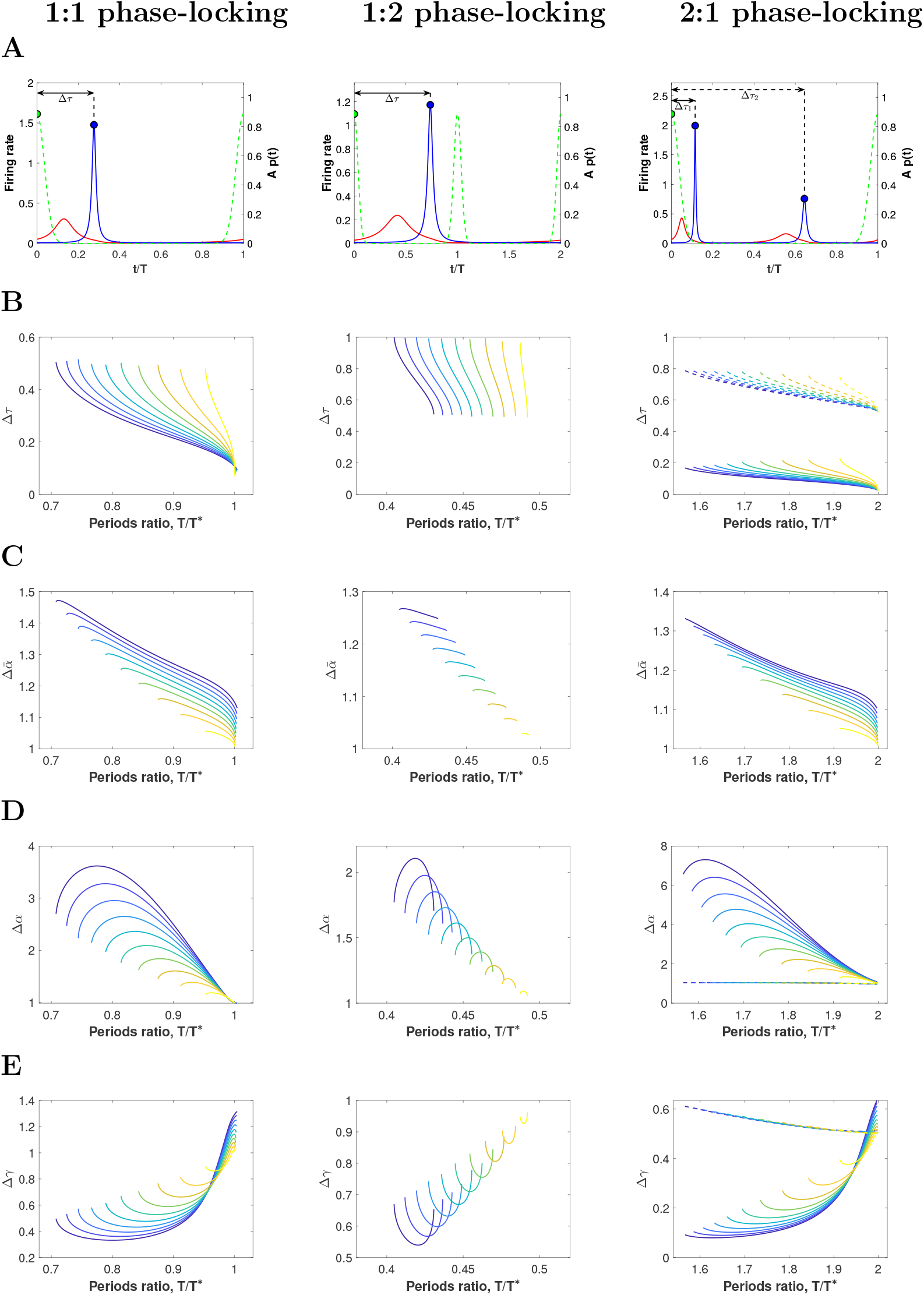
Input effectiveness for network entrained by inputs with high coherence. (A) Evolution of the firing rate variables *r_e_* (red) and *r_i_* (blue) and the von Mises perturbation for *κ* = 20 (dashed green) for a representative periodic orbit within the 1:1 (left), 1:2 (middle) and 2:1 (right) phase-locking regions. (B-E) Factors describing effective communication within these phase-locking regions for orbits calculated along (equidistant) sections *A* = ct of the corresponding Arnold tongues, indicated by the color of the curve (ranging from dark blue, *A* = 1, to yellow, *A* = 0.1, with increments of size 0.1). The factors are: (B) Δ*τ*, describing the timing between inhibition and input volleys (normalized by the input period *T*), (C) 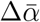, describing the rate change in the averaged firing rate, (D) Δ*α*, describing the rate change in the maximum of the (excitatory) firing rate, and (E) Δ*γ*, describing the rate change in E-volley half-width. See Methods. Additionally, the factors Δ*τ*, Δ*α* and Δ*γ* within the 2:1 Arnold tongue are also shown for the oscillator’s second peak (dashed lines).

We focus first on the 1:1 phase-locking region (see Figure 6 left). The entrainment presents different features inside the 1:1 Arnold tongue depending on the frequency of the external input. Indeed, on the right boundary of the Arnold tongue, corresponding to input frequencies close to the natural frequency of the gamma cycle (*T/T* ^*^ ≈ 1), the input volley reaches the target network when the I-cells are active (Δ*τ* ≈ 0). See Figure 6A and B left. Therefore, the overall firing rate of the E-population is barely affected by the external input but it grows with the amplitude *A* (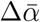 is close to 1 for small A, see Figure 6C left). However, as the input frequency increases (*T/T* ^*^ decreases away from 1) along a fixed amplitude inside the 1:1 Arnold tongue, we observe that the input volley tends to increase the time by which it precedes the I-population volley, thus arriving when the inhibition is at its minimum (Δ*τ* up to 0.5) and enhancing the response of the *E*-cells (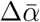 increases as *T/T* ^*^ decreases, see Figure 6C left). Thus, phase-locking is more effective for frequencies lower than the intrinsic network frequency (as observed for sinusoidal inputs in Section 2.3). See also Supplementary Figure S5.

Remarkably, it is interesting to explore the shape of the E-volley by means of the factors Δ*α* and Δ*γ* (Figures 6D and E). The non-monotonic shape of these factors inside the 1:1 Arnold tongue along a fixed amplitude is more pronounced than in the sinusoidal case (compare with Figure 4 left) and also for lower values of the input coherence *κ* (results not shown). Indeed, in the sinusoidal case, as soon as the input steps on inhibiton the 1:1 phase-locking is lost, while an input with higher coherence (*κ* = 20) can overcome the effect of inhibition more easily and therefore entrain the network for a wider range of frequencies. Thus, for high input frequencies (Δ*τ* ≈ 0.5 close to the left boundary) there is not enough time for the inhibition to decay before the arrival of the next input volley (decay time of inhibition is related to the synaptic time constant *τ_s_*), affecting the E-volley. For frequencies *T/T* ^*^ ≈ 1 (Δ*τ* ≈ 0 close to the right boundary), the factor Δ*α* ≈ 1 and Δ*γ* > 1 (see Figures 6D and E left). Therefore, the E-volley widens (i.e. the E-cells become less coherent) because the input arrives slightly later than the natural activation of the E-cells. Thus, it lengthens the time the E-cells are active without increasing significantly the firing rate due to the activation of inhibition (Δ*α* ≈ 1). We will see later in Section 2.6 that this effect is more pronounced in the case of oscillations close to the Hopf bifurcation.

In the 1:2 phase-locking region we observe a similar behavior as in the sinusoidal case, but the non-monotonic shape of the factors Δ*α* and Δ*γ* is again more pronounced (compare Figure 4 middle and Figure 6 middle). See also Supplementary Figure S6.

Finally, in the 2:1 phase-locking region, we find, for the amplitudes explored, two E/I-volleys per one of the input (see, for instance, Figure 6A right). In this case, as opposed to the sinusoidal case, where the input is wider and present in all the phases of the cycle, we observe that the input affects only one E/I-volleys (solid line in Figure 6 right), while the second E/I-volley reflects only the intrinsic dynamics of the network and it is barely modulated by the external input (Δ*α* and Δ*γ* barely vary for dashed line in Figure 6D and E right). For the affected E-volley, we observe the same properties as in the 1:1 phase-locking region, namely, the dependence of Δ*τ* on the input frequency and the non-monotonic shape of the factors Δ*α* and Δ*γ* (see solid curves in Figures 6 B, D and E right). See also Supplementary Figure S7.

### 2.5 Inputs in competition and selective communication

Thus far we have studied how a single periodic external input may entrain the network and how is the communication established between the emitting and receiving populations. In this section we want to explore how CTC theory implements selective communication. In selective communication the target network receives several stimuli from different sources, but responds only to one of them (the relevant stimulus), while ignoring the others [9].

To identify the relevant properties that are involved in selective communication, we probe how the phase-locked states described in the previous Section are affected by the presence of a distracting stimulus. In particular, we consider the case where the target network is entrained by a primary periodic input *p*_1_(*t*) of the form (17) (with frequency *f*_1_ = 1/*T*_1_ and coherence *κ*_1_ = 20) and we add a second periodic perturbation *p*_2_(*t*) of the form (17) for which we will vary the frequency *f*_2_ = 1/*T*_2_ and the coherence *κ*_2_ (see Figure 7A). We investigate whether the distractor prevents or not the target oscillatory network to follow the primary stimulus and how communication between populations is affected. This approach allows us to identify which stimulus is effective and which one is ignored and the underlying mechanisms for this selection.

**Figure 7:**
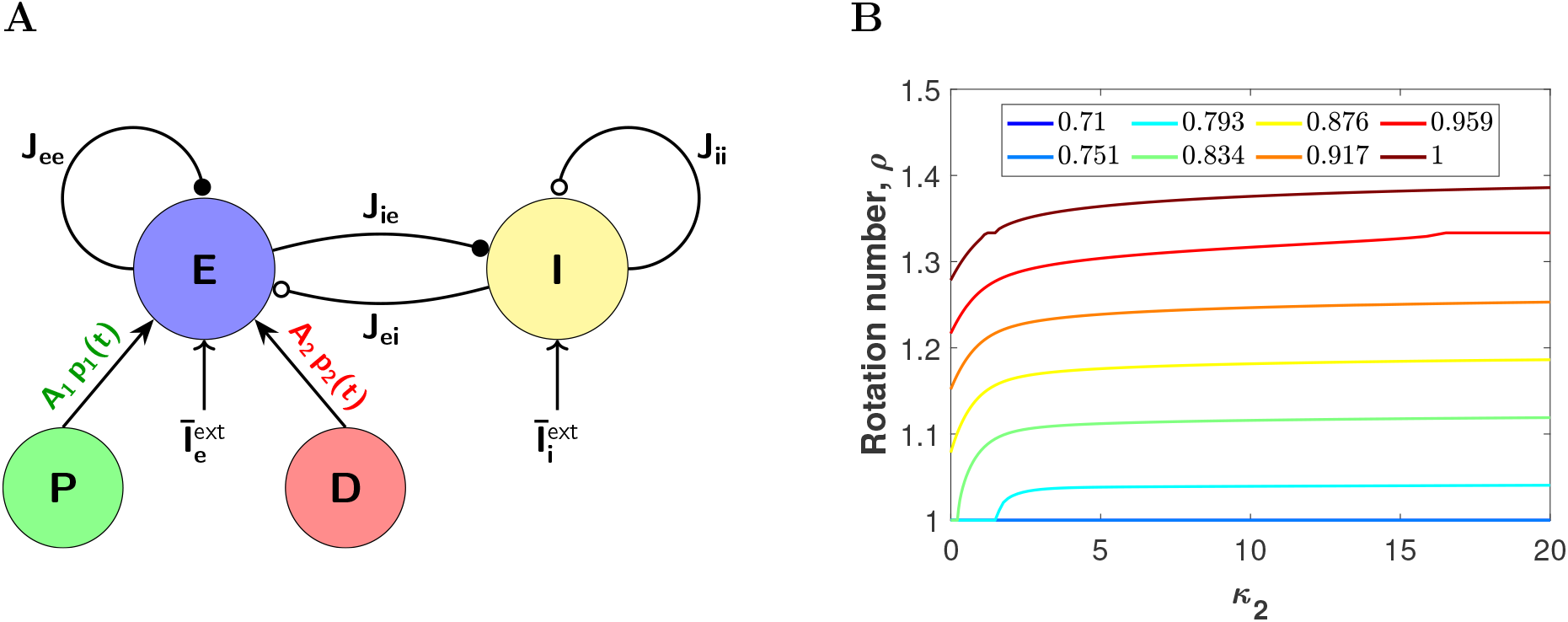
Effect of an identical frequency distractor in the network entrained by the primary stimulus. (A) Schematic representation of an E-I cortical neural network (PING interplay) receiving two oscillatory inputs from different sources: the primary input *A*_1_*p*_1_(*t*) (green circle) and the distractor one *A*_2_*p*_2_(*t*) (red circle). (B) Rotation numbers *ρ* of the stroboscopic map (9) for a perturbation consisting of a primary input and a distractor. Both inputs are modelled by means of a von Mises distribution, have the same amplitude factor *A*_1_ = *A*_2_ = 1 and the same period *T* = *T*_1_ = *T*_2_ but phase-shifted. The coherence for the primary is fixed *κ*_1_ = 20. We vary the the distractor coherence *κ*_2_ (x-axis) and the period *T*, so that the values *T/T* ^*^ (color legend) are equidistributed along the 1:1 plateau for *κ*_1_ = 20 (the oscillator and the primary stimulus support a 1:1 phase-locking relationship). If *ρ* = 1, the entrainment by the primary stimulus is preserved despite the presence of the distractor, otherwise, it breaks down.

#### 2.5.1 Two input streams of the same frequency

We first consider the case when the distractor has the same frequency than the primary input (i.e, *f*_2_ = *f*_1_ = 1/*T*_2_ = 1/*T*_1_ = 1/*T*), but phase-shifted by *T*/2 so that the two inputs are in anti-phase. We assume that the coherence of the primary is high and fixed at *κ*_1_ = 20 and we vary the coherence of the distractor *κ*_2_ and the period of both inputs *T*. The parameter is set at *A* = 1 and we consider only those values of *T* for which the primary input entrains the network, that is, the pair (*T/T* ^*^, *A*) lies inside the 1:1 Arnold tongue (see Figure 5D).

Since both inputs have the same frequency, the perturbed system is still *T*-periodic and therefore we can compute the rotation number *ρ* for the stroboscopic map (9) of the phase equation (8) with *p*(*t*) = *p*_1_(*t*) + *p*_2_(*t*). Recall that if *ρ* ≈ 1, then 1:1 phase-locking is preserved, whereas if *ρ* is away from 1, then the distracting stimulus manages to disrupt phase-locking.

The ability of a distractor to disrupt the entrainment by the primary stimulus depends mainly on the frequency relationship *T/T* ^*^ (see Figure 7B). Indeed, for lower values of *T/T* ^*^ (on the left hand side of the 1:1 Arnold tongue), the entrainment is more robust to perturbations by a distractor input. Remarkably, this effect is fairly independent of the coherence of the distractor *κ*_2_. For lower input frequencies the entrainment is not affected for all values of *κ*_2_ (the rotation number remains constant *ρ* = 1). As the input frequency decreases (*T/T* ^*^ increases), the entrainment by the primary is significantly affected by the distractor (*ρ* ≠ 1) and although the rotation number slightly increases away from 1 with *κ*_2_, it saturates at low values of *κ*_2_.

For the cases when the entrainment is not disrupted, we will see in Section 2.5.5 that the network may switch between the attended stimuli.

#### 2.5.2 Two input streams of different frequency

We now turn to study the effect of a distractor of different frequency than the primary that entrains the network. We fix the parameters *κ*_1_ and *T*_1_ of the primary input *p*_1_(*t*) so that there is a 1:1 phase-locking relationship between the primary and the oscillatory E-I network. Then, we add a second external input *p*_2_(*t*), for which we vary the coherence *κ*_2_ (between 0.1 and 20) and the period *T*_2_ (within the range 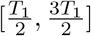). The distractor is phase-shifted so that both input volleys are as much apart as possible over a cycle (see Figures 8A,B and Methods for a precise definition). The amplitude for both inputs is set to *A* = 1.

**Figure 8:**
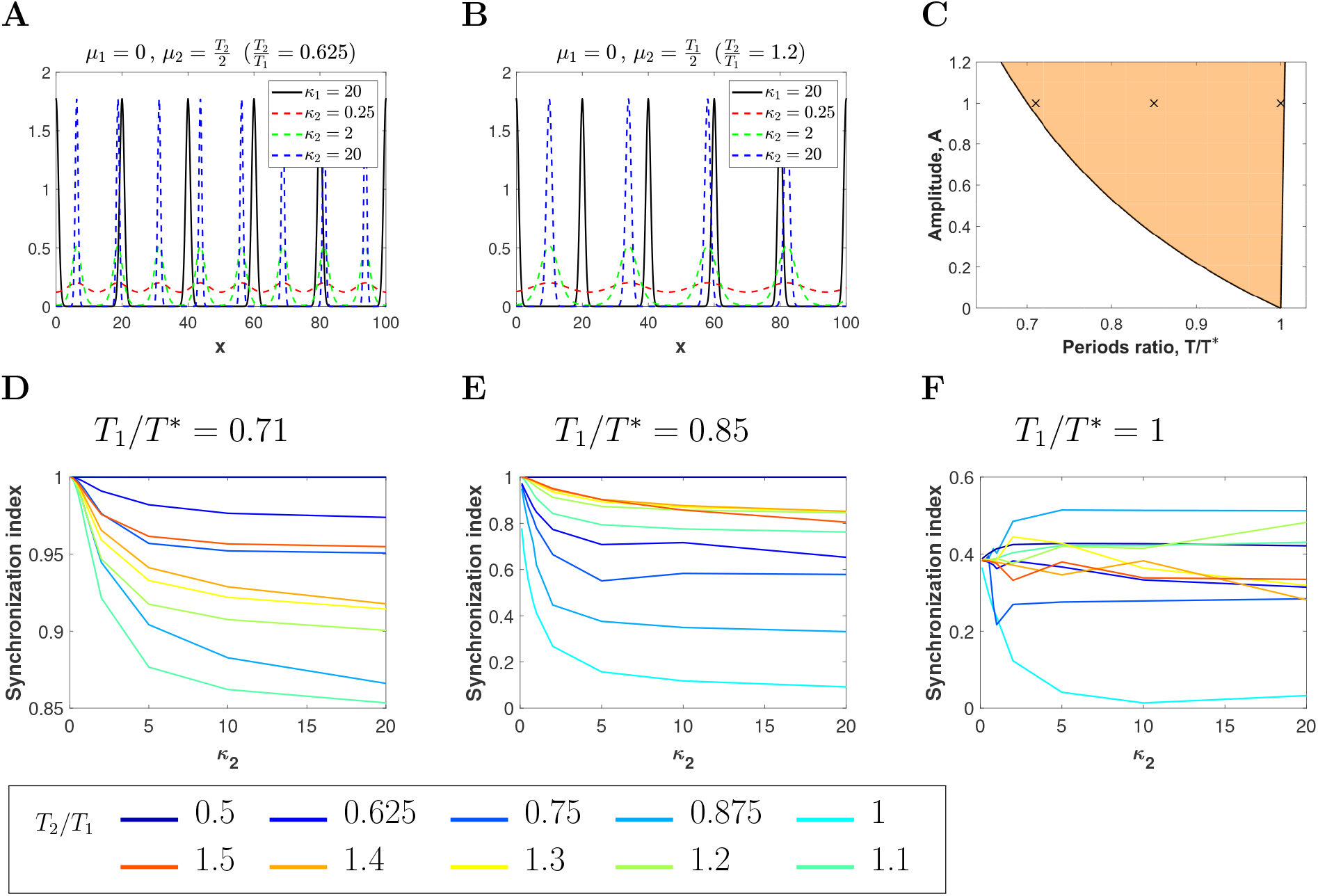
Effect of a non-identical distractor in network entrained by the primary stimulus. (*A*, *B*) Two von Mises inputs of different frequency phase-shifted according to the rule described in Methods. The black curve corresponds to the primary stimulus located about *μ*_1_ = 0 with *κ*_1_ = 20 (high coherence) and period *T*_1_ = 20. The dashed curves correspond to a distractor with different coherence values *κ*_2_: 20 (blue), 2 (green) and 0.25 (red). (A) *T*_2_/*T*_1_ < 1, the distractor is phase-shifted *T*_2_/2, (B) *T*_2_/*T*_1_ > 1, the distractor is phase-shifted *T*_1_/2. (C) Arnold tongue corresponding to 1 : 1 phase-locking between a single von Mises input with coherence *κ*_1_ = 20 (primary input) and the target network. We have selected 3 orbits along the section *A* = 1 (black crosses) corresponding to *T*_1_/*T* ^*^ = 0.71, 0.85 and 1 (left to right), for which we apply a distractor input. (D, E, *F*) Synchronization index *r* as a function of the coherence of the distractor *κ*_2_ (x-axis) for different values of the periods ratio between inputs, *T*_2_/*T*_1_ (color legend). The distractor frequency can be higher (cold colors) or lower (warm colors) than the primary’s frequency, being as much twice as fast (dark blue line) or 3/2 times slower (red line).

Notice that in this case the perturbed system (1)-(2) with *p*(*t*) = *p*_1_(*t*) + *p*_2_(*t*) is no longer *T*_1_-periodic, neither the phase equation (8). However, we still consider the time-*T*_1_ map of the phase equation, which provides a phase over the cycle at each time *T*_1_ (although the evolution of this map depends also on time). See Methods for more details. We compute the *synchronization index* (SI) *r*, also known as *vector strength*, for the phases of this map. The synchronization index is a measure of how clustered are the phases on a cycle [38, 39]. Thus, if *r* ≈ 1, the phases are all clustered around the same value, while if *r* ≈ 0 they are scattered around the circle. See supplementary Figure S8.

The frequency relation *T*_1_/*T* ^*^ has been fixed at three different points inside the 1:1 Arnold tongue for *κ*_1_ = 20 along the line *A* = 1: on the left (*T*_1_/*T* ^*^ = 0.71), in the middle (*T*_1_/*T* ^*^ = 0.85) and on the right-hand side of it (*T*_1_/*T* ^*^ = 1). See Figure 8C. In Figures 8D-F we show the SI as a function of the frequency relation between the primary input and the distractor (*f*_1_/*f*_2_ = *T*_2_/*T*_1_) and the coherence of the distractor *κ*_2_. We consider that the entrainment of the target network by the primary input is maintained despite the presence of the distractor when SI satisfies *r* > 0.8.

We observe that the entrainment is not disrupted by the presence of a distractor when the relation *T*_1_/*T* ^*^ lies near the leftmost edge of the 1:1 Arnold tongue (see Figure 8D for *T*_1_/*T* ^*^ = 0.71). The distractor can either oscillate faster or slower (within the range 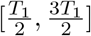) or it might even come in the form of sharper or broader pulses (vary *κ*_2_), but it does not prevent the primary input to entrain the oscillator. Notice that the SI remains above 0.85 for all values of *κ*_2_ and frequency relations *T*_2_/*T*_1_ (see Figure 8D).

When *T*_1_/*T* ^*^ lies in the center of the 1:1 Arnold tongue, faster distractors than the primary input (i.e. *T*_2_/*T*_1_ ≤ 1) cause the phase-locking breakdown since the SI falls below 0.8 (see cold color curves in Figure 8E), independently of the distractor coherence *κ*_2_, as long as it is above 1. However, slower distractors than the primary input (i.e. *T*_2_/*T*_1_ ≥ 1) do not disrupt phase-locking significantly. Indeed, the SI remains above 0.8 (see warm color curves in Fig 8E).

Finally, when *T*_1_/*T* ^*^ lies close to the right border of the 1:1 Arnold tongue, the distractor succeeds in completely disrupting the 1:1 phase-locking, irrespective of *κ*_2_ and the frequency relationship *T*_2_/*T*_1_ (see Figure 8F). Indeed, the SI for all values is below 0.5.

In conclusion, if the distractor is coherent enough (i.e. *κ*_2_ is sufficiently large), what determines the robustness of the entrainment is the frequency relationship *T*_1_/*T* ^*^. Indeed, the entrainment of the network by the primary input is much stronger (robust to distractors) when the primary has a much higher frequency than the natural gamma cycle. In this case, even the more coherent distractors cannot break the 1:1 phase-locking. As the primary input frequency decreases and approaches the natural gamma cycle frequency, the entrainment becomes weaker because, first, faster distractors can break the synchrony and finally, close to the right boundary, any distractor can do so.

We emphasize that when the primary input does no longer entrain the E-I network (due to the effect of the distractor), this does not necessarily mean that is being entrained by the distracting stimulus. We have computed the SI *r* for the phase map at time *T*_2_ and observed that *r* ≠ 1 (and far from 1) for most of the frequencies *f*_2_ = 1/*T*_2_ (results not shown). However, there are some cases in which this happens. Namely, when *T*_2_ is such that *T*_2_/*T* ^*^ lies close to the left branch of the Arnold tongue, then the role of the primary and the distractor are interchanged. For instance, when *T*_2_/*T*_1_ = 0.875 and *T*_1_/*T* ^*^ = 0.85 (see Figure 8E), and when *T*_2_/*T*_1_ = 0.875 (and 0.75) and *T*_1_/*T* ^*^ = 1 (see Figure 8F).

In the next section we will discuss the underlying mechanisms that explain the dependency of the entrainment robustness on *T*_1_/*T* ^*^.

#### 2.5.3 The role of inhibition

In the previous section, we observed that the entrainment is more robust for higher frequencies of the primary input. The explanation is related to the phase relationship that emerges between the primary input and the I-volley of the target network. First, notice that in the absence of the periodic primary stimulus, the E-I volley peaks are shifted apart by approx 0.18 phase units when 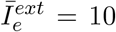 (see Figure 2D). We emphasize that this difference is maintained more or less constant when we add the entrainment either by a sinusoidal input or a von Mises type (results not shown).

Observe that for values of *T*_1_/*T* ^*^ close to left boundary of the 1:1 Arnold tongue, the factor Δ*τ* is close to 0.5 (see Figure 6B left), thus indicating that the I-volley arrives as much apart as possible (in phase) of the primary input volley. Therefore, inhibition is present when the distractor volleys arrive, thus suppressing their effects. In conclusion, inhibition shadows the distracting stimulus and phase-locking with the primary is not disrupted. See supplementary Figure S9 left.

On the contrary, if *T*_1_/*T* ^*^ is close to right boundary of the 1:1 Arnold tongue, Δ*τ* is close to 0 (around 0.1) (see Figure 6B), thus indicating that the peak of inhibition occurs close to the primary input. Therefore, inhibiton is over when the distractor volleys reach the E-cells of the target network, thus enhancing the effect of the distractor input onto the E-cell activity and giving the competitor the opportunity to disrupt phase-locking. See supplementary Figure S9 right.

For intermediate values of *T*_1_/*T* ^*^, only faster inputs disrupt phase-locking (*T*_2_/*T*_1_ < 1), because every time an input volley arrives under sufficiently low inhibition elicits (indirectly) an I-volley that shadows the effects of subsequent input volleys arriving close in time. Thus, faster distractor inputs are more frequent and are capable to elicit more I-volleys that shadow the effects of the primary volleys and disrupt phase-locking. See supplementary Figure S9 middle.

#### 2.5.4 Effects of the disruptor on the E-cell evoked response

We are not only interested in studying whether entrainment is preserved but also how the communication (i.e., E-cell evoked response) is affected by the presence of a distractor. More precisely, in those cases in which the entrainment by the primary input is maintained, we want to explore how the presence of a distractor affects the response of the E-cells of the target network to the primary input. A question that arises is whether the output of the E-I network reflects summation in some sense of the two inputs or whether one dominates over the other.

To quantify these effects, we compute the rate increase in the maximum of the firing rate of the E-cells *r_e_* and its mean firing rate over several cycles with respect to the non-entrained E-I network (factors Δ*α*^(0)^ and 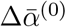, respectively, in Figure 9 top) and the entrained network by the primary (factors Δ*α*^(1)^ and 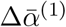, respectively, in Figure 9 bottom). Details of the computation of these factors are given in Methods (see equations (28) and (29)). Thus, magnitudes Δ*α*^(1)^ and 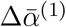 allow us to identify how the distractor affects the response of the E-cells to the primary, while Δ*α*^(0)^ and 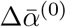, when compared with Δ*α*^(1)^ and 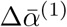, respectively, allow us to disentangle the contribution of the distractor and the primary to the response of the E-cells.

**Figure 9:**
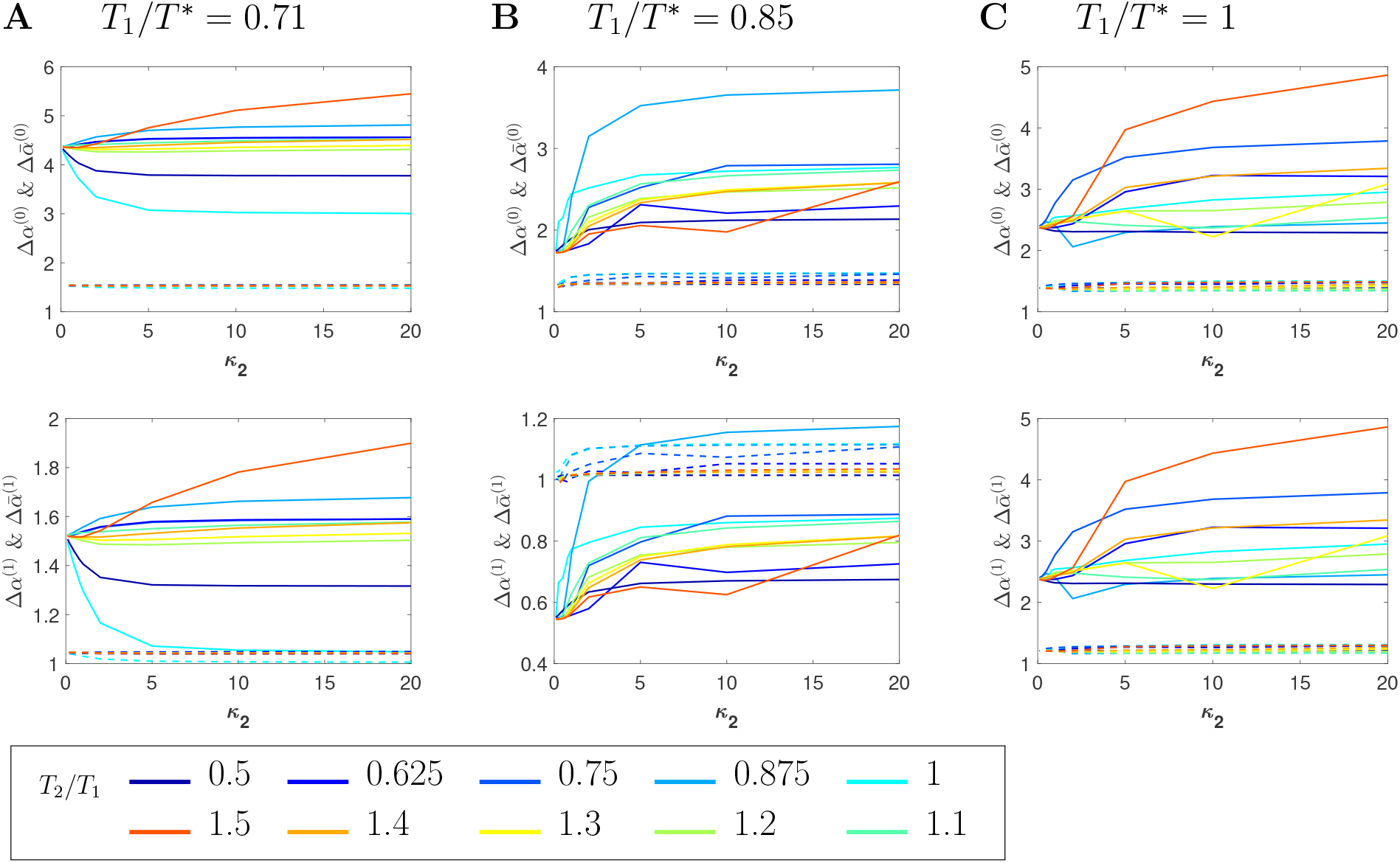
Effects of the disruptor on the E-cell evoked response. Factors measuring the impact of a (non-identical) von Mises distractor on the response of the E-cells of the E-I network when compared with (A) the natural gamma cycle (Δ*α*^(0)^, 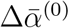, and (B) with the network entrained by the primary (Δ*α*^(1)^, 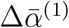, for the same parameter values as in Figure 8. Factors Δ*α*^(0,1)^ (solid lines) and 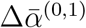 (dashed lines) measure the rate increase in the maximum of the excitatory firing rate *r_e_*(*t*) and the overall firing rate over a cycle, respectively. To compute factors Δ*α*^(0,1)^ we have averaged the maximum values of *r_e_*(*t*) over the first 20 peaks. Similarly, to compute factors 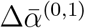 we have time-averaged *r_e_*(*t*) over the time interval 30max(*T*_1_, *T*_2_).

Remarkably, for values of *T*_1_/*T* ^*^ on the left-hand side of the 1:1 Arnold tongue, not only the entrainment is not affected by the distractor, but the maximum firing rate of the E-cells increases (Δ*α*^(1)^ > 1) when the distractor is present (see solid curves in Figure 9A). This happens because the I-volley is in anti-phase with the primary input volley. Thus, the windows for the disruptor to evoke a response are only open during the arrival times of the primary volley. Therefore, depending on the frequency relationship *T*_2_/*T*_1_, we have that, for some cycles, the input volley of the distractor and the primary input almost coincide, thus providing more concentrated input drive to the E-cells and evoking a higher response, while for other cycles, the distractor input volley reaches the E-cells when inhibition is present and the network does not respond to the distractor. See also supplementary Figures S9 left and S10A.

However, for values of *T*_1_/*T* ^*^ in the middle region of the Arnold tongue, the presence of a distractor decreases the evoked response of the E-cells to the primary input, even when the network is still entrained by the primary (for frequencies of the distractor *T*_2_/*T*_1_ > 1). Notice that values of Δ*α*^(1)^ are mainly below 1 (see solid curves in Figure 9B). The reason is again related to the timing of the I-volley. Recall that, as *p* increases, the time difference between the primary input volley and the I-volley shortens (see Figure 6B left), thus offering more temporal windows (away from the primary input volley) for the disruptor to elicit a response on the E-cells. This causes that for certain cycles the disruptor volley elicits a response ahead of the primary, thus affecting the entrainment and the overall response. See also supplementary Figures S9 middle and S10B. The case *T*_2_/*T*_1_ = 0.875 is different (Δ*α*^(1)^ > 1 here), because for these parameter values the primary and the distractor exchange their roles (as discussed in the previous section).

For the case *T*_1_/*T* ^*^ = 1, when we compare the response of the E-cells with no input (Δ*α*^(0)^) or with the primary input (Δ*α*^(1)^), we obtain exactly the same graph (see Figures 9C), because the primary input does not elicit any response on the E-cells, i.e. it is barely ignored (see Δ*α* ≈ 1 for *T*_1_/*T* ^*^ = 1 in Figure 6D). Adding a distractor increases significantly the response in the E-cells (values of Δ*α*^(0)^ and Δ*α*^(1)^ above 2.5 in Figure 9C solid curves), because now the target network responds to the distractor, thus enhancing the activity of the E-cells. See also supplementary Figures S9 right and S10C.

Notice that in all cases, the factors 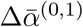, which measure the rate change of the mean firing rate of the E-cells, is above 1 but close to 1 (see dashed curves in Figure 9). Thus, the presence of a distractor does not increase significantly the overall firing rate of the E-cells, but it makes the response more or less coherent.

#### 2.5.5 Selective communication and switching between identical input streams

When the E-I network receives two identical input streams, we have identified a range of frequencies, for which the target network oscillates coherently (phase-locked) with both of them. Interestingly, the phase relationship that emerges causes that only one of the inputs affects the activity of the E-cells. Thus, the E-I network responds selectively to one of the inputs, while ignores the other. In this section, we want to explore the ability of the network to switch between the input that is attended. This has possible implications for selective attention [16].

Consider an oscillatory E-I neural network entrained by two identical input streams *p_i_*(*t*), *i* ∈ {1, 2} of the form (17) located in anti-phase (see, for instance, supplementary Figure S9C left). This situation occurs in the 1:2 Arnold tongue (see Figure 5D), and similarly, in the case of two identical inputs with a particular period relationship with the natural gamma cycle *T/T* ^*^ (those values such that the rotation number *ρ* = 1 in Figure 7B). Notice that in the latter case, the 1:1 phase-locking between the target network and the first input is not disrupted by the addition of a secondary one. The network is entrained such that the E-cells respond only to one of the input streams.

Mathematically, the stroboscopic map *P* in (9) of the phase equation (8) for the forced system with input *p*(*t*) = *p*_1_(*t*) +*p*_2_(*t*) has a stable periodic orbit of period two, or equivalently, the map *P* ^2^ has two stable fixed points. Therefore, depending on the initial condition, the trajectory approaches either one or the other, which translates to the network E-cells responding either to the first or the second input.

We study the effects of a brief/short stimulus (a pulse) to the entrained network. Brief stimuli could transiently lengthen or shorten the cycle period (i.e. phase shifting) and force the oscillator to respond to the other input, thus producing a change in the effective input. To quantify these effects, we consider the periodic solution for the perturbed 8-dimensional model (1)-(2) with *p*(*t*) = *p*_1_(*t*) + *p*_2_(*t*) and apply a square wave pulse of amplitude 3 and duration 2 ms (i.e. brief and short stimulus). In Figure 10 we show, for a particular case, the change in phase (blue curve) and in the effective input (black solid/dashed line), as a function of the phase *θ* of the entrained oscillator at which the pulse is applied. See Methods. Solid line denotes no change in effective input - the oscillator responds to the same input stream after the pulse - and dashed line denotes a change - the oscillator responds to the other input stream after the pulse. Note that the phase shift is positive almost for all phase, meaning that a pulse typically causes a phase advance (i.e. the E-volley occurs earlier). Notice that there is only a small range of phases (between 0.5 and 0.8) for which a square wave pulse produces a switch in the effective input. This interval corresponds to phases of the cycle for which the I-volley just passed and the next E-volley has not activated yet (see, for instance, Figure 10C). Notice that the phase shift is close to zero if the pulse arrives right when the I-volley activates.

**Figure 10:**
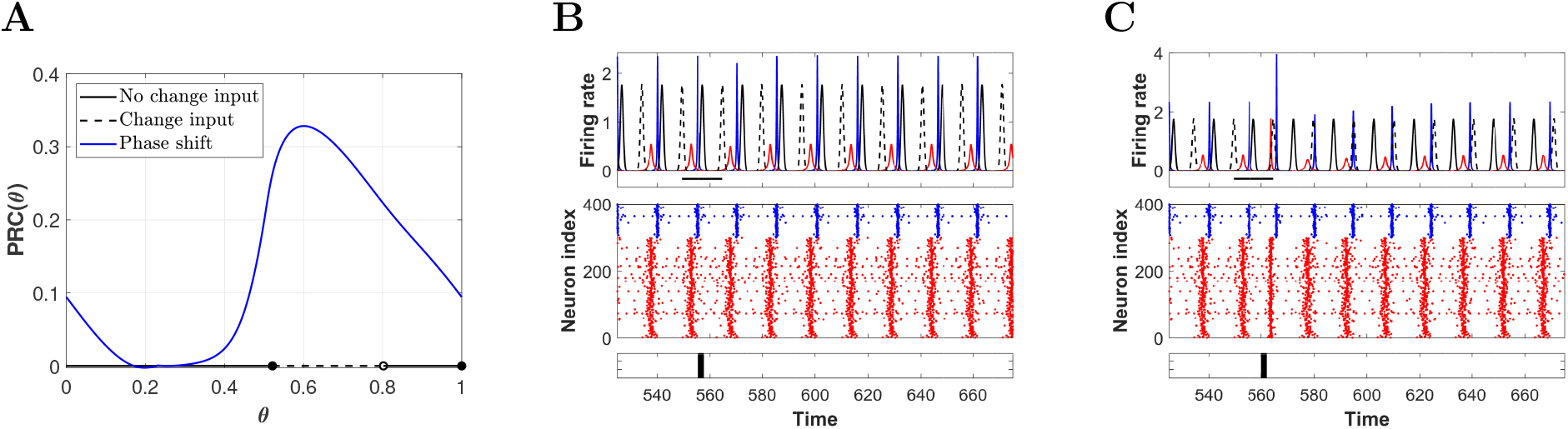
Selective communication and switching between attended stimulus. (A) Phase Response Curve (solid blue) obtained by applying square-wave perturbations of amplitude 3 and duration 2 at different phases *θ* of the periodic solution of system (1)-(2) receiving two identical inputs of von Mises type (*κ*_1,2_ = 20 and *T*_1,2_/*T* ^*^ = 0.73). The plot also shows if the (entrained) oscillator follows the same input before and after the pulse administration (black solid line) or if the attended input has changed (dashed black line). (B, C) Simulations of the full microscopic model showing the response of the network to a square-wave current delivered at two different phases of the cycle: (B) *θ* = 0.4, for which no switching between attended stimuli occurs, and (C) *θ* = 0.7, for which switching occurs. Each panel shows (from top to bottom), for a time interval of 150 ms, the two identical von Mises inputs in antiphase (solid and dashed black) with the mean firing rates of the E-cells (red) and I-cells (blue) of the network, the corresponding raster plot and the time at which the square-wave pulse is applied. We have integrated the full network of QIF neurons for 1000 ms. At time 200 ms we apply the two inputs of von Mises type. The square-wave pulse is applied at time (23 + *θ*)*T* + 200 ms.

We simulate the full microscopic model to support and illustrate these results. A pulse perturbation of the full microscopic model at phase 0.4 does not entail a change in the effective input (oscillator still responds to the dash-lined input after the pulse is applied in Figure 10B), while at phase 0.7 it does cause a change in the effective input (oscillator responds to the solid-lined input after the pulse is applied in Figure 10C), as predicted by the reduced model.

### 2.6 Oscillations close to the Hopf bifurcation

Thus far we have observed that the timing and shape of the I-volley in the gamma cycle plays a relevant role in implementing several aspects of the CTC theory. To emphasize better this role, we explore how CTC theory implements in a different type of gamma cycles. More precisely, we consider oscillations arising from the PING mechanism but closer to the Hopf bifurcation curve in the parameter space (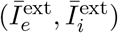) (see Figure 2A). Thus, for parameters as in (6) and tonic currents to the E and I-cells 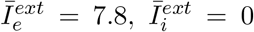, the system (1)-(2) presents oscillations at a frequency 1/*T* ^*^ = 1/33.433 ≈ 30Hz (lower gamma range) of the form shown in Figure 11A. Notice that E/I-volley are wider and lower for this oscillation than the one considered along this paper with 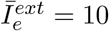 (compare with Figure 3A).

**Figure 11:**
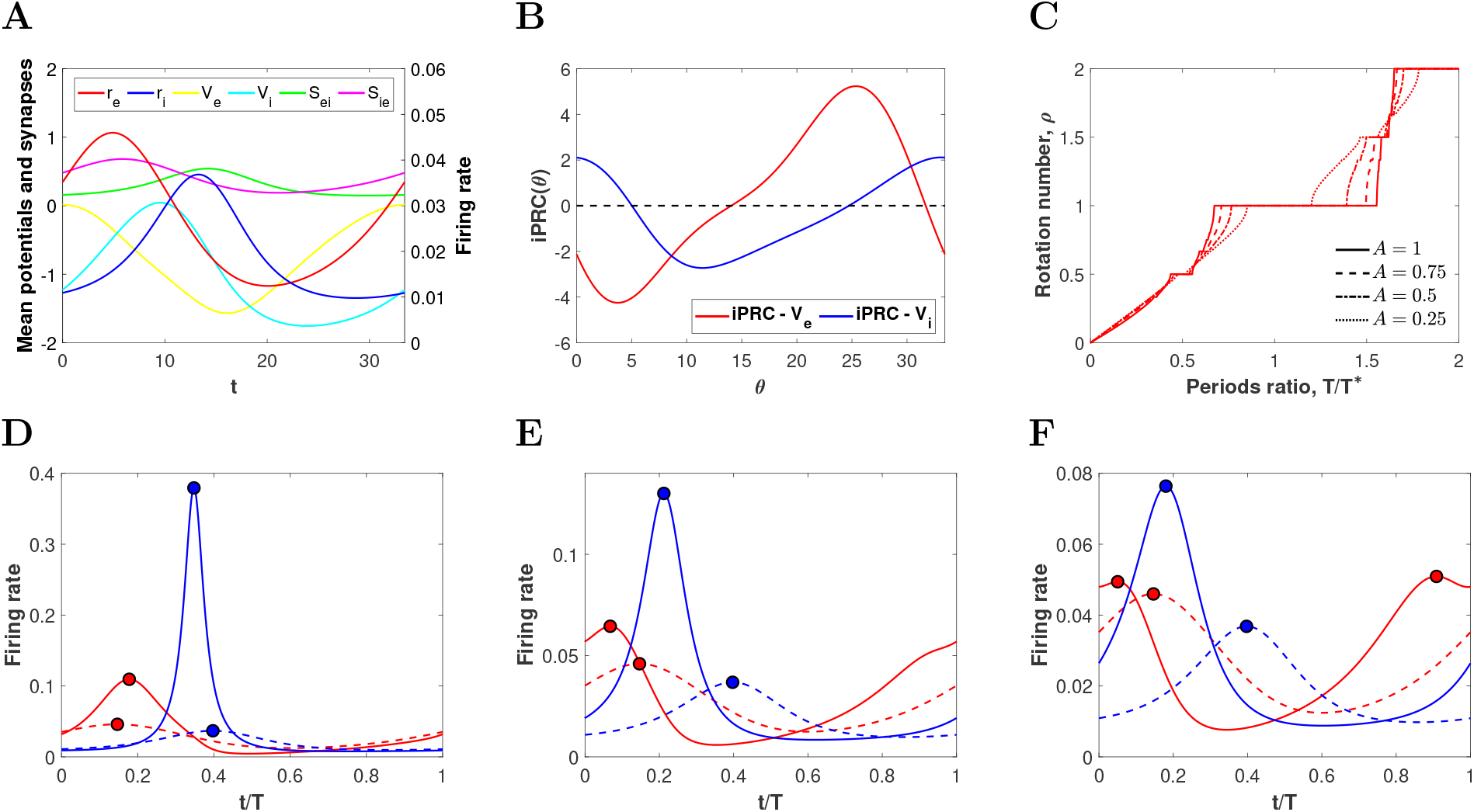
PING oscillations close to a Hopf bifurcation. (A) Temporal evolution of firing rate, mean membrane potential and synapses variables over a cycle of a PING oscillation corresponding to external current 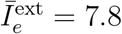 (close to the Hopf bifurcation curve in Figure 2A). (B) Infinitesimal Phase Response Curve (iPRC) of the cycle in Panel *A* for perturbations in the direction of the variables *V_e_* and *V_i_* (red and blue curves, respectively). Note that the iPRC 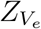 is both positive and negative. (C) Rotation numbers of the stroboscopic map (9) for a von Mises input (16) with coherence *κ* = 20 applied in the direction of *V_e_*, as a function of the ratio between the intrinsic period of the E-I network *T* ^*^ and the input period *T* and different amplitude factors *A*. (D, E, *F*) Time evolution of the mean firing rates of the E-cells (red) and I-cells (blue) along the unperturbed (dashed curves) periodic orbit for the system (1)-(2) and perturbed (solid curves) with the high-coherent von Mises input with *A* = 0.25 and relative frequency *T/T* ^*^: (D) 0.8, (E) 1 and (F) 1.05. The period of the oscillators has been normalized to 1. Notice that the point *T/T* ^*^ = 0.8 does not lie on the 1:1 plateau for *A* = 0.25 predicted by the phase reduction.

Moreover, we also compute the iPRC for this oscillator (see Figure 11B). The iPRC takes both positive and negative values, as opposed to the oscillator with 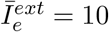 (compare with Figure 3B). This means that this oscillator may be entrained by either faster or slower periodic inputs. This is the typical behavior for oscillations close to a Hopf bifurcation [31].

We perturbed this oscillator with a periodic input with high coherence, *κ* = 20 (see equation (17)) and applied the phase reduction method described in Section 2.2. The rotation numbers for the stroboscopic map (9) as a function of the relative frequency (*T/T* ^*^) are plotted in Figure 11C for different amplitude values. The visible plateaus, corresponding to rational rotation numbers (*ρ* = *p/q*), are centered about the point *T/T* ^*^ = *p/q* and extend towards both sides. Therefore, the oscillator can be entrained by faster (i.e. *T/T* ^*^ < *p/q*) and slower inputs (i.e. *T/T* ^*^ > *p/q*), as expected from the iPRC shape. Notice that the largest plateaus correspond to 1:1 and 2:1 phase-locking (*ρ* = 1 and *ρ* = 2, respectively) and they grow as the amplitude *A* increases. Moreover, the transition from *ρ* = 1 and *ρ* = 2 becomes sharper (see the devil’s staircase for *A* = 1 in Figure 11C). This result is similar to the one observed in [24] for the Wilson-Cowan model.

Since the rotation numbers in Figure 11C have been computed using the phase reduction approach, the phase-locking predictions may not be accurate for the perturbed 8-dimensional mean-field model (1)-(2), specially when the amplitude increases. Indeed, we observe that the actual phase-locked solutions do not extend as far to the right (towards lower input frequencies) but extend further to the left (towards higher input frequencies) than the predicted by the phase reduction. Thus, for *A* = 0.25, the phase reduction identifies 1:1 phase-locking for *T/T* ^*^ ∈ (0.85, 1.2), but the actual computation finds synchronized solutions only for *T/T* ^*^ ∈ (0.757, 1.075).

Even if 1:1 phase-locking for low input frequencies does not extend as far to the right as predicted, we still find entrained solutions for inputs with frequencies lower than the network intrinsic frequency (*T/T* ^*^ > 1). It is interesting to explore how is the response in the target network for low input frequencies, compared to those with high input frequencies. We look for periodic solutions of the perturbed meanfield model (1)-(2) along the 1:1 phase-locking plateau for *A* = 0.25. For *T/T* ^*^ = 0.8 (high input frequency) the entrained oscillator shows a significant increase of the firing rate activity of both E and I-cells (compare solid and dashed lines in Figure 11D). As the frequency of the input becomes more similar to the natural gamma cycle, the excitatory firing rate presents an inflection point right before reaching the maximum (see Figure 11E for *T/T* ^*^ = 1), that becomes a small second peak (see Figure 11F for *T/T* ^*^ = 1.05). This is because the E-cells activate due to both, the intrinsic properties of the network and the input volley, the latter arriving shortly after the E-cells start to activate but before they can trigger inhibition. Thus, the inputs with low frequencies contribute to widen the E-volley, but without increasing its maximum significantly (the E-cells become less coherent). For larger values of *T/T* ^*^ we do not find periodic solutions anymore.

To better identify the aspects related to CTC theory, we compute the factors Δ*τ* and 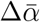 (as well as Δ*α* and Δ*γ*) introduced in Section 2.3 to measure the timing of the inhibition and the changes in the excitatory firing rate due to the perturbation, respectively. We have computed such magnitudes for the solutions in the 1:1 phase-locking region for amplitudes *A* = 0.25 and *A* = 0.5 (yellow and blue curves, respectively, in Figure 12). Clearly the factor Δ*τ* indicates that the input volley precedes inhibition (0 < Δ*τ* < 0.5) and the temporal distance Δ*τ* reduces as the input frequency decreases (*T/T* ^*^ increases) (see Figure 12A). Accordingly, the factor 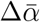 measuring the increase in the overall firing rate also reduces as *T/T* ^*^ increases (see Figure 12B). Interestingly, for values *T/T* ^*^ above 1, the increase in the overall excitatory firing rate (Figure 12B) and maximum of the E-volley (Figure 12C) is weak (values of 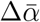 and Δ*α* close to 1), but the width of the E-volley increases (Δ*γ* above 1 in Figure 12D), thus causing a decrease in the firing coherence of the E-cells of the target network.

**Figure 12:**
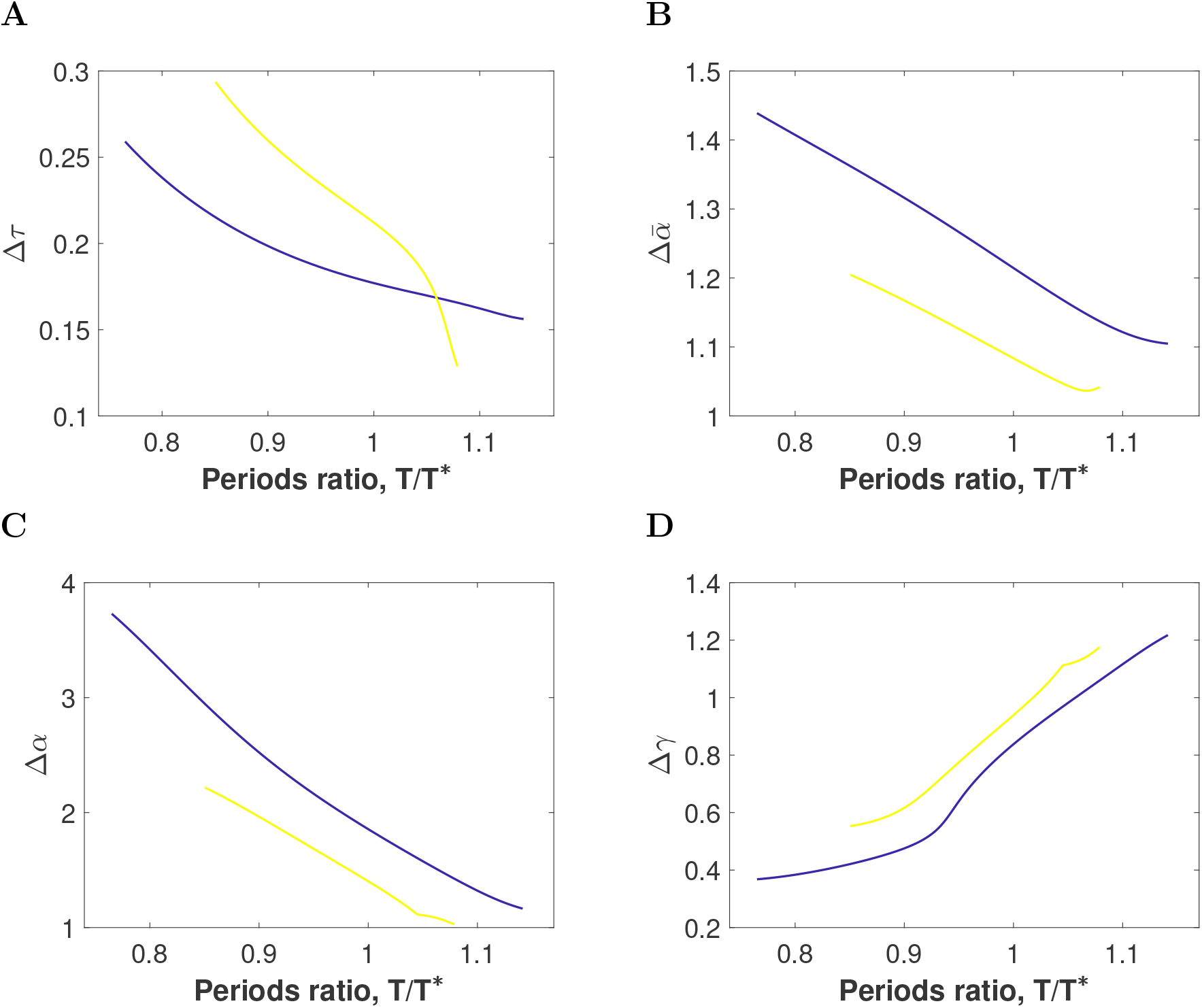
Input effectiveness for entrained network close to a Hopf bifurcation. Factors (A) Δ*τ*, (B) 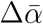, (C) Δ*α* and (D) Δ*γ* describing effective communication within the 1:1 phase-locking regions for orbits calculated along the amplitudes *A* = 0.25 (yellow curves) and *A* = 0.5 (blue curves). Notice that these factors are computed in smaller intervals than those predicted by computing the rotation number using the phase reduction (see Figure 11C). The non-smooth point observed in the Δ*α* curve around *T/T* ^*^ 1.05 for *A* = 0.25 is due to the presence of two peaks in the excitatory firing rate (only one for the I-cells, see Figure 11F) and a switch in the maximum between them.

## 3 Discussion

In this paper we have studied several hypothesis of the CTC theory, namely, the conditions for optimal phase-locking and selective communication. Our approach considers a network of E-I cells oscillating in the gamma range, whose macroscopic dynamics can be exactly described by a low-dimensional mean-field model. In the communication setting of CTC theory, the E-I network receives external time-periodic input, which models the effect of the emitting population. We vary the amplitude, frequency and coherence (how concentrated are inputs in a cycle) of the external input and explore the conditions under which phase-locking between the input and the target population occur and how the phase relationship that emerges contributes to communication. We then focus on the response of the E-I network when it receives input streams in the gamma frequency range from two different populations. We observe that inputs with higher coherence can entrain the network for a wider range of frequencies. Besides, for higher input frequencies the inhibitory volley does not interfere with the input volley and the response of the network (in terms of increase in firing rate) is stronger than for input frequencies similar to the intrinsic network gamma cycle. Moreover, the entrainment by higher input frequencies is more robust to perturbations.

We stress that the use of mean field models of low dimension allows for mathematical analysis. In particular, we have identified semi-analytically (with numerical bifurcation analysis) the phase-locking regions (Arnold tongues) in the parameter space, when we vary the frequency, amplitude and coherence of the external input. Our mathematical analysis confirms that, in general, phase-locking occurs through a tilted Arnold tongue (bent towards higher input frequencies) as observed in computational studies of spiking neurons [18]. Indeed, only faster inputs can entrain the network because the input volley can only evoke an excitatory response and end the gamma cycle if it arrives under sufficiently low inhibition [37].

Besides, we have provided more details about the entrainment by higher frequency inputs. Namely, we have observed that increasing the amplitude of the stimulus enlarges and also shifts the range of frequencies that entrain the network towards lower values, while increasing the coherence enlarges the range without shifting it (see Figures 3D and 5D). Thus, our results confirm analytically previous computational studies on spiking networks claiming that E-cells in the PING target network show preferential phase-locking to periodic inputs with higher coherence. This phenomenon was referred as ‘coherence filtering’ in [37].

It is hypothesized that E-I networks generating gamma rhythms automatically produce an optimal phase relationship because when a sufficiently strong excitatory pulse is delivered to the network, it generally elicits and I-volley (indirectly in the case of PING). If the frequency of the forcer is approximately the frequency of the receiver, the inhibition from the I-volley wears off just when the next input volley is due to arrive. Thus, the entrainment automatically sets up an optimal phase relationship for CTC [37, 20]. In this paper, we have gone beyond this result and provided a quantitative description of this optimality in terms of the effects in the E-cell evoked response (maximum and coherence) of the target network, as well as the robustness of the entrainment to distractors. In our study we have found that the phase difference is optimal in the sense that the input precedes inhibition, but the phase difference is not the same for all frequencies that are capable to entrain the E-I network (in line with [18]), and this has consequences for communication. Indeed, those inputs with higher frequencies precede the I-volley by far, thus causing a strong amplification of the activity of the E-cells in the target network. In contrast, those inputs with frequencies similar to the receiver almost step on the I-volley, thus having a weaker effect onto the activity of the E-cells. Interestingly, this finding is independent of the coherence of the input. Thus, synchrony of the source network does not necessarily produce higher firing in the target network.

In our study, we have also explored the effects of input frequency and coherence in selective communication, the phenomenon in which a target network entrains and responds to only one of several input streams. This mechanism is conjectured to implement selective attention, the process in which subjects focus on a particular object of the environment while ignoring the others. In this context, experimental studies in visual cortex have shown that when two stimuli are presented in the receptive field of a neuron, it can respond to either input depending on whether attention is directed to [14, 40]. Indeed, selective attention is associated with an increase in coherence of spiking and in spectral power of oscillations in the gamma frequency band [14, 41, 42].

Previous modeling studies [22, 37] have observed that higher coherence and frequency lend a competitive advantage to an oscillatory presynaptic neuronal population to entrain a target oscillating network. Here, we have gone a step further since we have explored how these two properties interact with each other and, interestingly, we have observed that coherence affects little, compared to frequency. Indeed, the higher the frequency of the primary stimulus that entrains the network the more robust to desynchronization by disruptors. The distractor (regardless its coherence) can easily break the synchronization of the network with a low frequency primary input, but not for a high frequency one. When the primary input has a much higher frequency than the natural gamma cycle (leftmost regions of the 1:1 Arnold tongue), the position of the I-volley in the cycle makes the entrainment more robust since most of the distractor volleys arrive when inhibitory cells are active. In contrast, for input frequencies similar to the natural gamma cycle (rightmost regions of the 1:1 Arnold tongue), the position of the I-volley lies close to the primary volleys, thus some of the distractor volleys arrive when the I-cells are silent, eliciting a response of the E-cells that disrupts synchronization with the primary, even for low coherence distractors. This result is in agreement with the experimental results in the visual cortex [40], where V1 sites processing relevant stimuli have its gamma peak frequency higher than the irrelevant V1 sites and higher than the target sites in V4.

As a novelty, we have observed that effective communication is not affected by the distractor when the entrainment by the primary is preserved, that is, the disruptor does not have a negative effect on the E-cells evoked response.

We have also explored how phase shifting can act as a mechanism for stimulus selection. In line with [25], an appropriate optogenetic pulse applied at the right phase can cause switching between the attended inputs. Our results show that the appropriate phases are after the inactivation of the I-volley but before the activation of the E-volley.

The mathematical formalism based on the phase equation used herein requires that the inputs received by the target network are weak [31, 33]. We emphasize though that the predictions on synchronized states obtained from the phase reduction in Figures 3, 5 and 11, were validated computing the corresponding periodic orbit for the the reduced mean-field model (1)-(2). However, we emphasize that in our previous study on Wilson-Cowan equations [24] we did not assume that inputs were weak, and we detected regions in the parameter space showing bistability between synchronous and asynchronous regimes, as well as between different types of entrainment. The method used herein was not designed though to detect bistability in the forced network, therefore the existence of bistability or other dynamical features in the strong coupling regime remains a challenging topic for further studies. Techniques based on the phase-amplitude description might be useful to tackle larger amplitude perturbations [43, 44, 45].

Changes in excitability are generated by the interaction between excitation and inhibition, and the frequency and phase relationship between *E* and *I* volleys can be tuned by means of changes in tonic drive. In our previous work with Wilson-Cowan equations [24], the dynamics of the oscillating firing rates of the E and I populations resembled those of the E-I network considered herein when 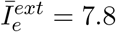. Namely, the E and I volleys are less coherent and they spread along the whole cycle (see Figure 11A). Moreover, the oscillation frequency of the network falls in the lower range of gamma oscillations (around 30Hz). Notice that, in this case, the oscillations are close (in parameter space) to a Hopf bifurcation (see Figure 2A) and the iPRC is of Type II (taking both positive and negative values). Thus, as opposed to the case with 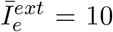, the target oscillating network can synchronize to both faster and slower incoming periodic inputs (see Figure 11B). Noticeably, in this case the prediction by the phase reduction is less accurate for the slowest inputs. This case makes clear the different effect of lower and higher frequency inputs than the natural gamma cycle in the activity of the target network. Indeed, higher input frequencies tend to increase both the firing rate and coherence of the network while lower ones tend to make it less coherent (see Figure 12).

We acknowledge the fact that gamma-band rhythms observed in experimental data do not show regular oscillatory behavior, but they are rather irregular and episodic [46, 47]. In [21] the authors consider an E-I network showing irregular and episodic gamma rhythms and observe that this assumption enables a target population to be correlated to two independent sources that differ either in frequency or in phase. Notice that in this case correlations are used to measure entrainment and synchronization between neural populations. We believe that the conclusions obtained for regular oscillations can though provide the substrate to explain several mechanistic phenomena in the irregular case. For instance, we have observed that it is possible to switch the entrainment between two different sources by means of pulses delivered at the proper phases of the cycle. This could be an interesting subject of research for future work.

In this paper we have mainly focused on the interactions between two neuronal groups oscillating in the gamma range, but we have not discussed the interaction of gamma with other rhythms, namely beta (13-30 Hz), alpha (8-13 Hz) and theta (4-8 Hz) [35, 37]. The interplay between different frequency rhythms may definitely have an important role for communication and cognition [37, 9, 48, 49, 50] and the methods discussed herein may contribute to unveil possible mechanisms. Namely, a thorough study of the solutions emerging in the p:1 phase-locking regions, where the input is *p* times slower than the intrinsic oscillatory frequency. We have studied the 2:1 phase-locking regions and observed that the slow input modulates the E/I-volleys of the gamma cycle, in a similar or different way depending on the frequency relationship between both rhythms *T/T* ^*^ and the input coherence (see Figure 4 right and Figure 6 right).

Finally, we want to acknowledge that CTC proposes other hypothesis that we have not included in this study. Thus, we leave for future work the study of the effects of inputs onto the inhibitory cells (see for instance [51]), as they have been proposed as a mechanism for phase-shifting in models of cortical networks [25]. Preliminary results have shown phase-locking for both lower and higher input frequencies. We have also mentioned in the introduction that gamma oscillations can be generated by means of an ING (Interneuron Network Gamma) mechanism [4]. Exploratory results have shown no remarkable differences regarding phase-locking with the ING mechanism, but a thorough study is required to draw conclusions. Finally, our setting can be extended to networks of two populations to explore bidirectional communication [52, 10, 29, 53] and its main differences with the unidirectional communication setting explored in this paper.

## 4 Methods

### 4.1 Neuron models

We consider a population of *N* neurons with all-to-all coupling, subdivided into a population of *N_e_* excitatory and *N_i_* inhibitory cells, whose individual voltage dynamics are modelled by a Quadratic Integrate-and-Fire (QIF) model [54], i.e.,

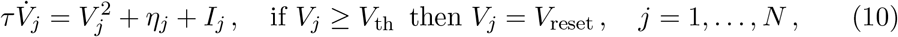

where *V_j_* is the voltage of the *j*-th neuron, *τ* is the time constant, *η_j_* is a constant bias current and *I_j_* is an input current accounting for external (time-dependent) stimuli and the excitatory/inhibitory synapses of the network. The threshold and reset voltages are taken *V*_th_ = −*V*_reset_ → ∞, (in numerical simulations, we will set them to *V*_th_ = −*V*_reset_ = 500). Heterogeneity in the system is introduced by assuming that the parameters *η_j_* are distributed according to a Lorentzian distribution with half-width Δ and centered at 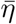,

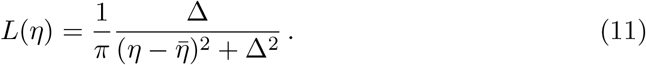

The input current *I_j_* consists of a common external input for all neurons and synaptic current due to recurrent connexions in the circuit. For the excitatory ensemble, the input is expressed as

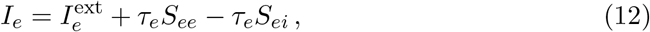

while for the inhibitory one as

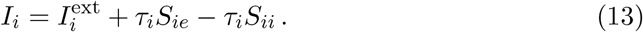

We have omitted the subindex *j* (labeling each neuron in the population) because all the neurons in the same population receive the same input current. The subscript indicates whether it corresponds to the excitatory (E) or the inhibitory (I) population. The variable *S_ab_* models the synaptic current from the presynaptic population *b* to the postsynaptic population *a*, where *a*, *b* ∈ {*e*, *i*}. The dynamics of the synaptic currents are described by a linear differential equation of the form

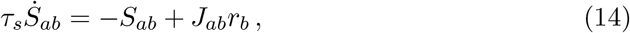

where *a*, *b* ∈ {*e*, *i*}. Here, *τ_s_* is the synaptic time constant, *J_ab_* the synaptic strength from population *b* to population *a*, and *r_b_* the firing rate of population *b*, which is computed as

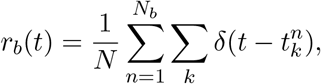

where *δ*(*t*) is the Dirac delta function and 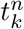 are the firing times of neuron n.

In [26, 29], one can find the details of the derivation of the reduced mean field model (1)-(2) that provides an exact description (in the thermodynamic limit) of the macroscopic quantities of the spiking model described above, namely, the mean firing rate *r_a_* and membrane potential *V_a_*.

### 4.2 Inputs

We consider different types of periodic functions *p*(*t*) to model the excitatory input drive to the E-cells 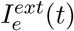 (see equation (5)). We consider first a sinusoidal input, i.e.

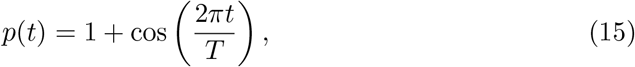

where *T* is the period of the perturbation.

We also consider a non-sinusoidal input, for which we can modulate its coherence, that is, how synchronized are the spikes of the external drive over a cycle. To do so, we use the von Mises probability density function (also known as the circular normal distribution) defined on the circle for the angle *θ* ∈ [−*π*, *π*] as:

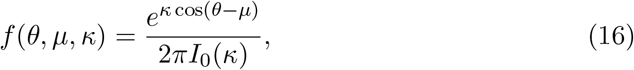

where *I*_0_(*x*) is the modified Bessel function of order 0. The parameter *μ* is the mean and *κ* is the temporal coherence (*κ* = 0 corresponds to a uniform distribution and as *κ* increases the distribution becomes more concentrated about the angle *μ*). See Figure 5A.

The *T*-periodic input *p*(*t*) is defined as

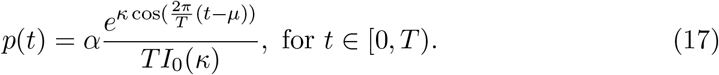

where *α* = *T*/2*π* so that the amplitude of the stimulus is independent of *T* and only varies with *κ* (see Figure 5A). When only one input is present, we fix *μ* = 0 and we vary the concentration factor *κ* and the period *T*.

We also consider the case when the E-population receives two input streams of the form (17), one referred to as the primary stimulus and denoted by *p*_1_(*t*), and the other as the distractor and denoted by *p*_2_(*t*). The mean, temporal coherence and period of the *i*-th input will be denoted *μ_i_*, **κ*_i_* and *T_i_*, respectively.

When both inputs have the same period (*T*_1_ = *T*_2_ = *T*) we place the second input to be in antiphase with respect to the first one, that is *μ*_2_ = *T*/2. However, when the periods are different the last criterion is not well-defined. Therefore, we implement the following rule: we place the peak of the input with largest period between two consecutive peaks of the input with shortest period. That is, if *T*_2_ ≥ *T*_1_, then 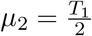, whereas if 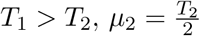.

### 4.3 Phase reduction

For an oscillating E-I network, the 8-dimensional system (1)-(2) has a hyperbolic attracting limit cycle Γ of period *T* ^*^. The limit cycle can be parameterized by the phase variable *θ*, such that *θ*(*t*) = *t* + *θ*_0_ ( mod *T* ^*^) and there exists a function *γ* : [0, *T* ^*^) → ℝ^8^ such that

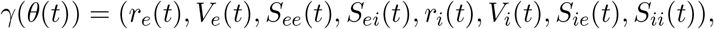

parameterizes the periodic orbit Γ.

We apply an external periodic input *Ap*(*t*) to the limit cycle Γ in the direction given by the vector ***v*** ∈ ℝ^8^. If the external periodic drive *Ap*(*t*) is weak, i.e. |*A*| << 1, we can reduce the study of the dynamics of the perturbed system to a single equation for the phase variable, given by

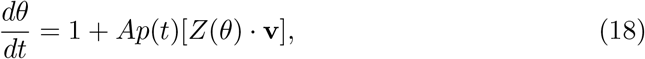

where the function *Z* : [0, *T* ^*^) → ℝ^8^ is the infinitesimal Phase Response Curve (iPRC). The iPRC measures the oscillator’s phase shift due to an infinitesimal perturbation applied at different phases of the cycle [31]. Notice that *Z*(*θ*) is a vector of 8 components, thus, the *i*-th component (*Z_i_*) corresponds to the phase shift due to an infinitesimal perturbation applied in the direction of the *i*-th variable (i.e. 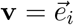). It is well known that the iPRC is the periodic solution of the so-called Adjoint Equation [31], given by

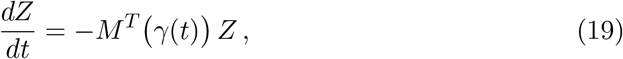

subject to the normalization condition

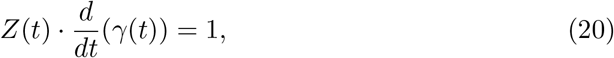

where the matrix *M γ*(*t*) is the linearization of the system (1)-(2) around the limit cycle Γ.

### 4.4 Rotation number

The existence of periodic points for the stroboscopic map (9) defined on the circle is given by a very well known result in the context of circle maps [36, 55]. Let 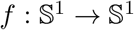 be a map on the circle, orientation preserving and regular enough, then the *lift* of *f* is a continuous function 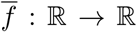 that satisfies 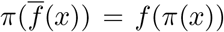, where *π*(*x*) = *x* mod 1 (i.e. the phase on the circle). Then, the *rotation number ρ* of the map *f* is defined as

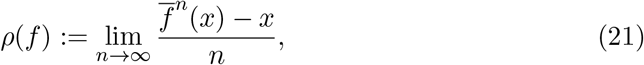

for any *x* ∈ ℝ. Importantly, the limit above always exists and does not depend on the initial point *x*, nor the lift 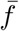 chosen.

There are two important results to characterize the dynamics of the map *f* using the rotation number. If *ρ* is rational (*ρ* = *p/q* with *p*, *q* ∈ ℕ), then there exists a *q*-periodic point of the map *f*, that is, a solution of *f*^*q*^(*θ*) = *θ* mod 1. If, by contrast, *ρ* is irrational, then the orbits of *f* fill densely the circle and there are no periodic points.

We compute the rotation number for the stroboscopic map (9) defined on the circle. We use the numerical procedure described in [56] to compute the rotation number of the stroboscopic map (9) in Figures 3C, 5B, 5C and 11C. We have used *N* = 750 iterations, and the error is of order 10^−6^.

### 4.5 Arnold tongues

The boundaries of the Arnold tongues correspond to the locus of a saddle node bifurcation [57] of the stroboscopic map (9) in the (*T*, *A*)-parameter space.

The saddle node bifurcation curves, corresponding to the boundaries of the *p* : *q* Arnold tongue, can be numerically computed by identifying the points (*θ*, *T*, *A*) such that the following two conditions hold:

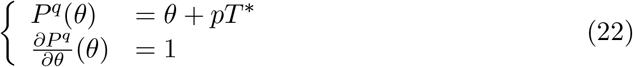

Notice that, although not specified, the stroboscopic map *P* depends on the parameters *T* and *A*. Moreover, since we take modulus *T* ^*^, we can omit the term *pT* ^*^ in the equation above.

The equations above describe a 1-dimensional curve in ℝ^3^, that can be computed using a continuation method. The continuation method is a numerical procedure by which one can find a curve in ℝ^n^ defined implicitly by a set of (nonlinear) equations *G*(*w*) = 0, where *G* : ℝ^n+1^ → ℝ^n^ is regular enough. Notice that this corresponds to a system with *n* nonlinear equations with *n* + 1 variables. The method to find the curve combines a tangent-like approximation as in the Keller’s (pseudo-arclength) method and a modified Newton method (based on a minimization problem with a restriction) to refine it. More precisely, starting from an initial solution *w*_0_ lying on the curve, *G*(*w*_0_) = 0, the method consists in making a prediction for the next point on the curve by moving along the tangent line to the curve at the point *w*_0_. Then, we correct the approximate point successively by means of a modified Newton method. Since the number of variables is higher than the number of equations, we need to impose the additional condition that the norm of the correction must be a minimum. We solve this problem using Lagrange multipliers. The details of the method can be found in [58, 56].

In our case, the problem consists of three variables (*θ* and the parameters *T* and *A*, after allowing them to evolve according to 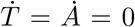) and two equations (22). Let *w* = (*θ*, *T*, *A*) be the unknowns and 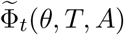 be the extended flow associated to the phase equation (8) when *T* and *A* areated as variables, and 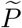 the associated stroboscopic map. Therefore, we define the function *G* as

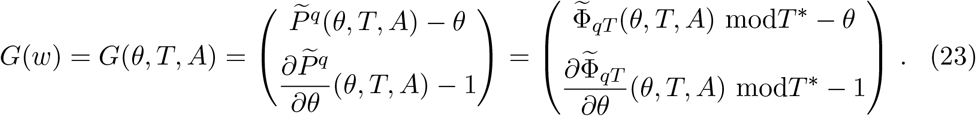

To start the method we must provide an initial seed. When the amplitude is zero and the forcing period satisfies a rational relation 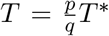, then any phase *θ* is a solution of *G*(*w*) = 0 (i.e. any phase provides a *p* : *q phase-locked* state of the forced oscillator). However, this point does not serve as initial seed because the tangent is not well-defined here. Instead, we consider *A* ≠ 0 and a period *T* at the edge of the plateau *ρ* = *p/q* of the Devil’s staircase and we look for a stable fixed point *θ* of the associated stroboscopic map by means of a bisection method.

Notice that the Newton method requires to compute the differential of the map *P*, which will be computed using variational equations [58]. Moreover, variational equations involve the computation of the derivative of the iPRC, which is only known numerically for a discrete set of points. Taking advantage of the periodic behaviour of the iPRC, we have used the FFT algorithm to compute the derivatives efficiently.

### 4.6 Measures of phase-locking

We present four magnitudes that will contribute to the description of the phase relationship between the input and the entrained network and determine how is the communication between neuronal populations in the CTC context. These magnitudes are computed for *qT*-periodic orbits of the perturbed system (1)-(2) with a given function *p*(*t*) of period *T*, whose existence is guaranteed in the *p* : *q* phase-locking regions.

The magnitude Δ*τ* computes the normalized time difference between the maximum of the I-volley *r_i_* and the maximum of the external perturbation *p(t)*, that is,

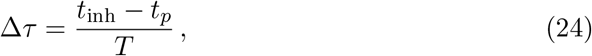

where *t*_inh_ and *t_p_* denote the time where the maximum of *r_i_*(*t*) and the perturbation *p*(*t*) are achieved over a cycle of period *T*, respectively. Notice that if Δ*τ* lies in the interval [0, 0.5), then inhibition follows the input, which may allow an increase of the activity of the E-cells of the target network. In contrast, if Δ*τ* lies in the interval [0.5, 1), then inhibition precedes the input and the latter may be ignored, bringing on a small or negligible effect onto the activity of the E-cells of the target population.

In the 1:2 phase-locking region, where there are two peaks of the perturbation *p*(*t*) per one of the inhibitory firing rate *r_i_*(*t*), Δ*τ* measures the distance between the peak of *r_i_*(*t*) and the preceding input peak (see, for instance, Figure 6A middle). Notice that Δ*τ* lies in the interval [0, 1). If Δ*τ* > 0 but close to 0, the inhibition immediately follows the first peak and precedes by far the second one. If Δ*τ* < 1 but close to 1, inhibition immediately precedes the second peak and follows by far the first one. For intermediate values of Δ*τ* ≈ 0.5, the I-volley is equidistant from both input peaks.

The effects of the perturbation onto the activity of the E-cells of the target population can be measured as well. We define 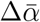 as the ratio between the time-average (over a single period [0, *qT*)) of the excitatory activity for the perturbed case (A ≠ 0), 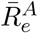, and the time-average of the excitatory activity for the unperturbed case (A = 0), 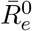, in the following way

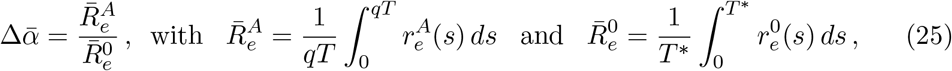

being 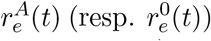 the first component of the periodic solution for the perturbed (resp. unperturbed) system.

Notice that if 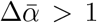, the external oscillatory input increases the response of the excitatory receiving population. Therefore, the larger 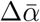, the more effective is the communication between the input source and the target oscillatory population. However, the effects of the perturbation onto the receiving population might be also described in terms of enhancement of synchronization in the receiving population. To measure so, we compute the quantities Δ*α* measuring the changes in the maximum and Δ*γ* measuring changes in the half-width.

Following [24], we define Δ*α* as the ratio between the maximum of the excitatory activity *r_e_*(*t*) for the perturbed case (A ≠ 0), 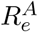, and the maximum of the excitatory activity *r_e_*(*t*) for the unperturbed case (A = 0), 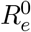, that is,

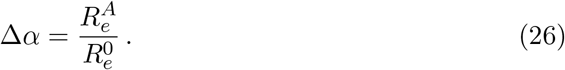

Notice that if Δ*α* > 1, the external oscillatory input increases the maximal response of the excitatory receiving population. Indeed, we may think that changes in the peak height indicate whether the external forcing synchronized (Δ*α* > 1) or desynchronized (Δ*α* < 1) the spikes of the target circuit.

The factor Δ*γ* provides the rate change of the half-width of the E-volley due to the external stimulus. It is defined as

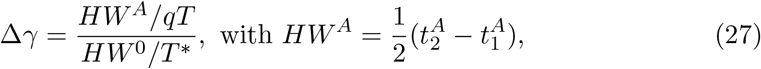

where 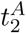 and 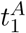 correspond to the two times 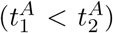 at which the firing rate is equal to half of its maximum 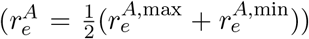. Notice that since we will deal with oscillators of different periods, we must normalize the time difference by its corresponding period.

To compute the factors Δ*α*^(0,1)^ and 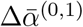 in Figure 9 we proceed in the following way. Recall that, in the presence of the distractor, *r_e_*(*t*) is not necessarily periodic. Therefore, we integrate the system (1)-(2) perturbed by two input streams (*p_i_*(*t*), *i* ∈ 1, 2 with amplitude *A_i_* and period *T_i_*) enough time to include at least 20 peaks of the excitatory firing rate activity that we denote as 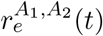, and we take the mean of the maxima. Then, we divide the result by the maximum of the excitatory firing rate over a period *T* ^*^ if the network is not entrained and *T*_1_ when the network is entrained by the primary input. This provides the factor Δ*α*^(0,1)^. More precisely,

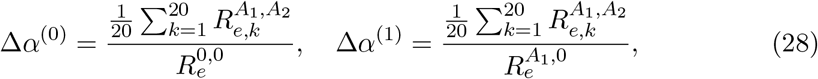

where 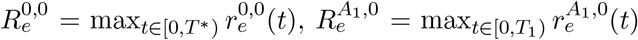 and 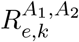 is the k-th local maxima of 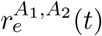 in the interval 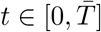, with 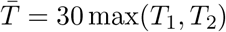.

Similarly, we compute 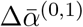 as

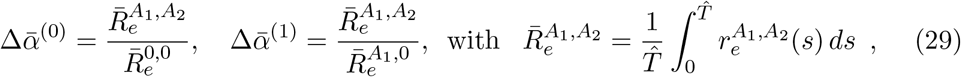

where 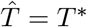 for 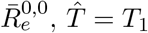 for 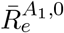 and 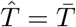 otherwise.

### 4.7 Vector strength

When we add a distractor of different frequency than the primary input, the time-*T*_1_ map of the solution of the phase equation (8) does not satisfy the conditions to compute the rotation number, since the second input (the distractor) is not *T*_1_-periodic. In this case, we still regard the phase in intervals of *T*_1_ time units (although they are now time-dependent) in order to analyse whether the distractor disturbs the entrainment and by which amount. To do so, we compute the *synchronization index* (SI) [38], also known as vector strength or Kuramoto order parameter [39]. It is a measure of how clustered are the events over a cycle. To compute the SI one associates to each event a vector on the unit circle with a phase angle and computes the mean vector. The Si is given by the length of the mean vector. That is,

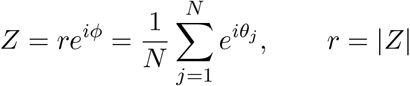

Notice that perfect clustering is obtained when *r* = 1, whereas if phases are scattered around the circle, then *r* ≈ 0.

### 4.8 Computation of the phase shift of the numerical PRC

The Phase Response Curve in Figure 10A provides the phase shift due to the application of a square wave current to the oscillation emerging in the E-I network (1)-(2) when entrained by two identical inputs in anti-phase. It is computed as follows. For every phase *θ* ∈ [0, 1) of the oscillator, we apply a square wave current pulse (amplitude 3, duration 2) and compute 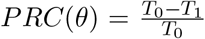 the period of the first cycle after the pulse application and *T*_0_ the period of the entrained oscillator.

The plot also shows, as a function of the phase *θ*, if the (entrained) oscillator switches between attended stimulus. To detect whether there is a change in the effective input, we perform the following steps:

1. Before applying the pulse (and once having converged to the periodic orbit) we measure, in a single cycle, the time distance from the maximum of the excitatory firing rate, 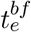, to the maximum of each input, 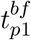 and 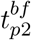. If 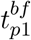 or 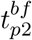 are larger than 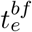 we consider the preceding input peak. Finally, we take the difference between these two distances, that is,

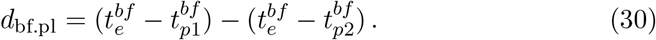
2. After applying the pulse, we integrate the whole perturbed system for long enough time (we have used 30 periods *T*) so that the transient effects have washed out. Then, we measure again the time distance from the peak of the excitatory firing rate to the peak of each input as in step 1. We take again the difference between these two time distances, keeping the same orientation as in step 1:

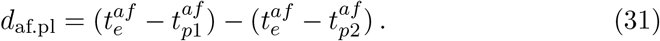

A change of sign between *d*_bf.pl_ and *d*_af.pl_ (i.e. sgn(*d*_bf.pl_ · *d*_af.pl_) = −1) will determine a change in the oscillator’s effective input.

### 4.9 Numerical methods

Equations for the 8-dimensional system (1)-(2) and for the phase equation (8) were integrated numerically in Matlab using an explicit Runge-Kutta method of order 4-5 (ode45) with an absolute tolerance ranging beween order 10^−12^ and 10^−16^.

Equations of the microscopic model consisting of *N_e_* = *N_i_* = 5000 QIF neurons were integrated using an Euler method with time step Δt = 10^−4^. There is a refractory period for each neuron of duration *T_ref_* = 2 · *τ*_*e,i*_/*V*_*th*_, where *V*_*th*_ = 500 is the voltage threshold and *τ*_*e,i*_ = 10 as in [59].

The bifurcation diagram in Figure 2A was computed using the numerical bifurcation analysis toolbox MatCont (a Matlab continuation package) [60, 61].

We used Matlab to analyse and plot the data.

Matlab codes have been released and are available at https://github.com/david-reyner/Matlab-Code.

## 5 Acknowledgements

Work produced with the support of a 2019 Leonardo Grant for Researchers and Cultural Creators, BBVA Foundation. The Foundation takes no responsibility for the opinions, statements and contents of this project, which are entirely the responsibility of its authors. GH acknowledges the RyC grant RYC-2014-15866. We acknowledge the use of the UPC Dynamical Systems group’s cluster for research computing (see https://dynamicalsystems.upc.edu/en/computing/).

## Supporting Information

**Figure S1:**
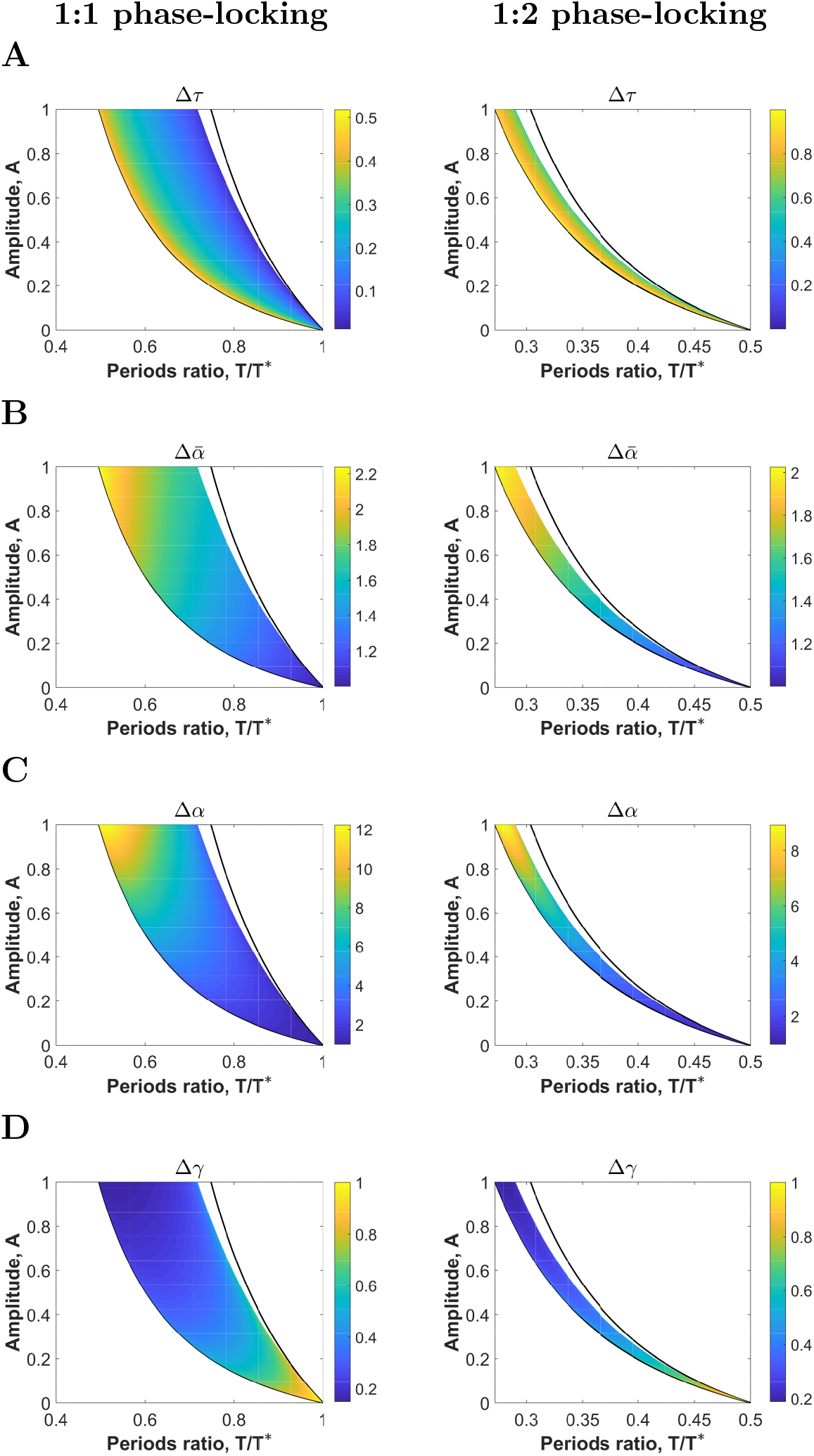
Input effectiveness for network entrained by sinusoidal inputs. Factors measuring effective communication for orbits within the 1:1 phase-locking (top) and 1:2 phase-locking (bottom) regions up to *A* = 1: (A) Δ*τ*, describing the timing between inhibition and input volleys (normalized by the input period *T*), (B) 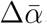, describing the rate change in the averaged firing rate, (C) Δ*α*, describing the rate change in the maximum of the (excitatory) firing rate, and (D) Δ*γ*, describing the rate change in E-volley half-width. See Methods. The factors have been computed only for values (*T/T* ^*^, *A*) for which the algorithm converges to a periodic orbit of system (1)-(2) perturbed with a sinusoidal input (colored region). Black solid lines correspond to the boundaries of the phase-locking regions predicted under the phase reduction hypothesis.

**Figure S2:**
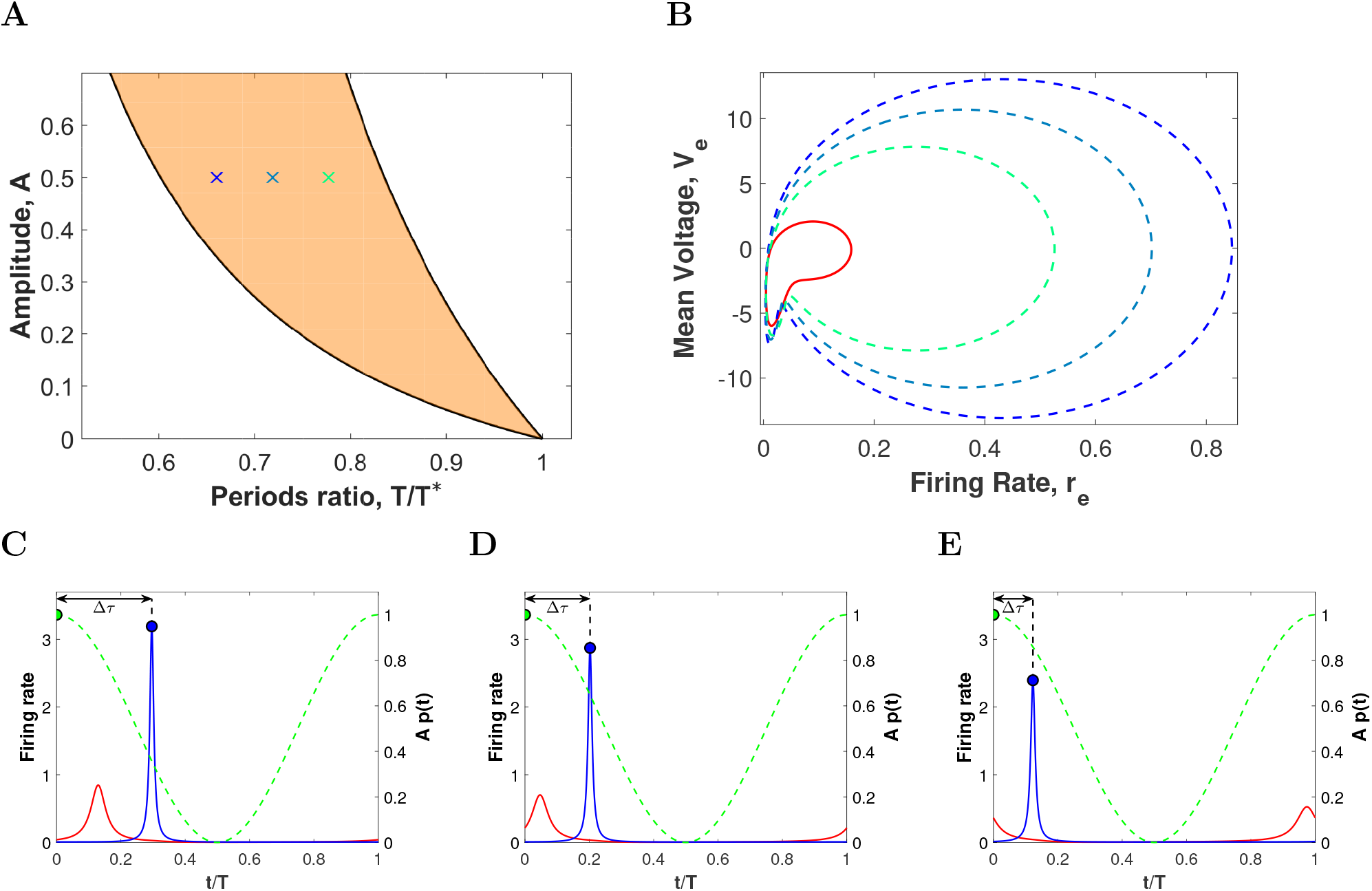
Representative periodic orbits within the 1:1 phase-locking region for sinusoidal inputs. (A) 1:1 phase-locking region between the PING oscillation in Figure 3A and a sinusoidal input (15) varying amplitude factor *A* and period ratio *T/T* ^*^. (B) Projection onto the (*r_e_*, *V_e_*)-plane of 3 representative periodic orbits corresponding to the crosses in panel *A* (dashed curves with same color as the crosses). Solid red line corresponds to the projection of the unperturbed limit cycle. (C-E) Evolution of the firing rate variables *r_e_* (red) and *r_i_* (blue) and the sinusoidal input *Ap*(*t*) (green dashed line) over a single period *T* for the three representative orbits: (C) left-hand side of the Arnold tongue (blue cross), (D) center of the Arnold tongue (turquoise cross) and (C) right hand side of the Arnold tongue (green cross). Blue and green circles indicate the maximum of the I-cells firing rate and the input, respectively. The magnitude Δ*τ* has been pointed out with a double-head arrow.

**Figure S3:**
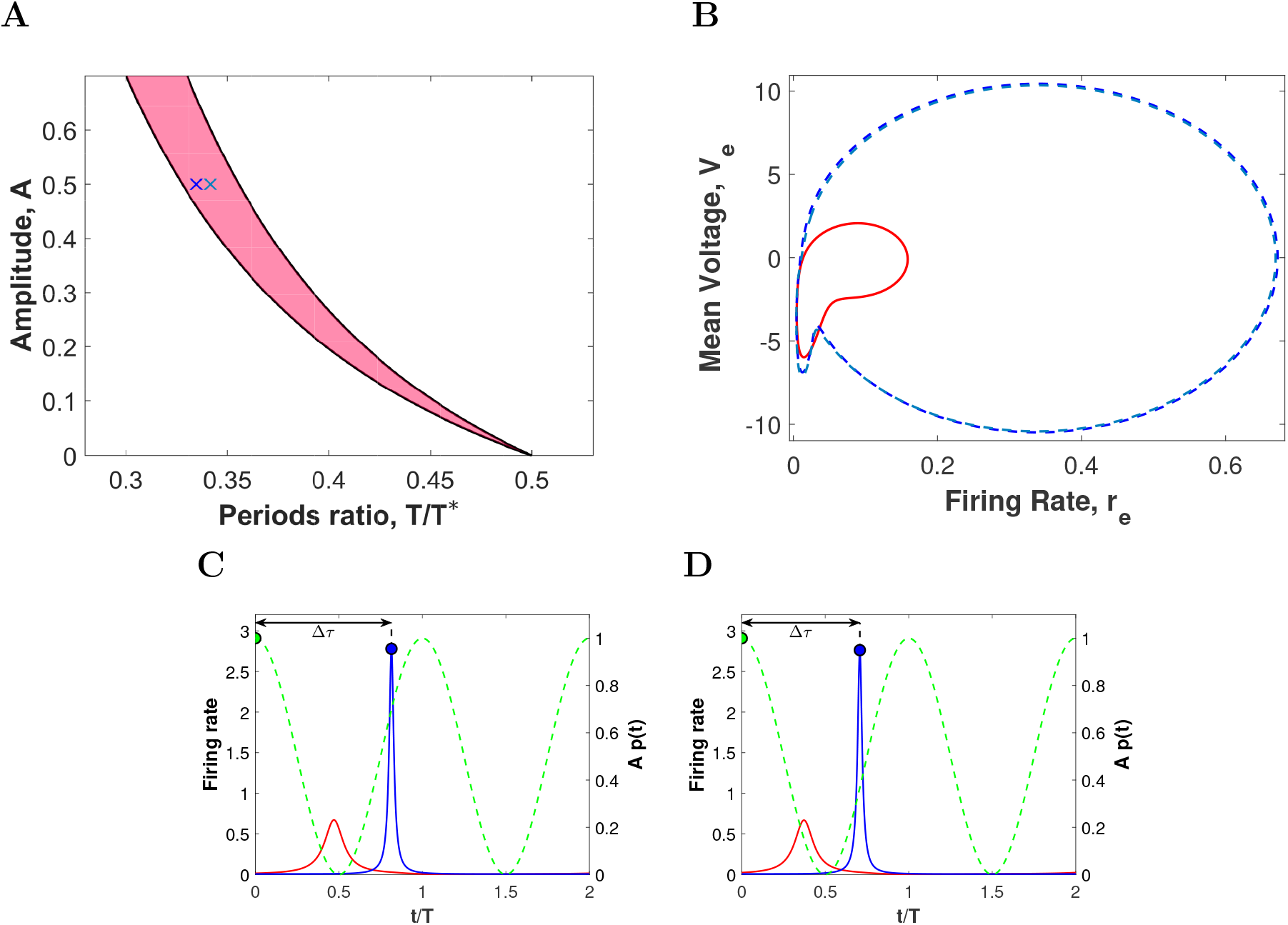
Representative periodic orbits within the 1:2 phase-locking region for sinusoidal inputs. (A) 1:2 phase-locking region between the PING oscillation in Figure 3A and a sinusoidal input (15) varying amplitude factor *A* and period ratio *T/T* ^*^. (B) Projection onto the (*r_e_*, *V_e_*)-plane of 2 representative periodic orbits corresponding to the crosses in panel *A* (dashed curves with same color as the crosses). Solid red line corresponds to the projection of the unperturbed limit cycle. (C-D) Evolution of the firing rate variables *r_e_* (red) and *r_i_* (blue) and the sinusoidal input *Ap*(*t*) (green dashed line) over two periods *T* for the two representative orbits: (C) left-hand side of the Arnold tongue (blue cross), (D) center of the Arnold tongue (turquoise cross). Blue and green circles indicate the maximum of the I-cells firing rate and the input, respectively. The magnitude Δ*τ* has been pointed out with a double-head arrow.

**Figure S4:**
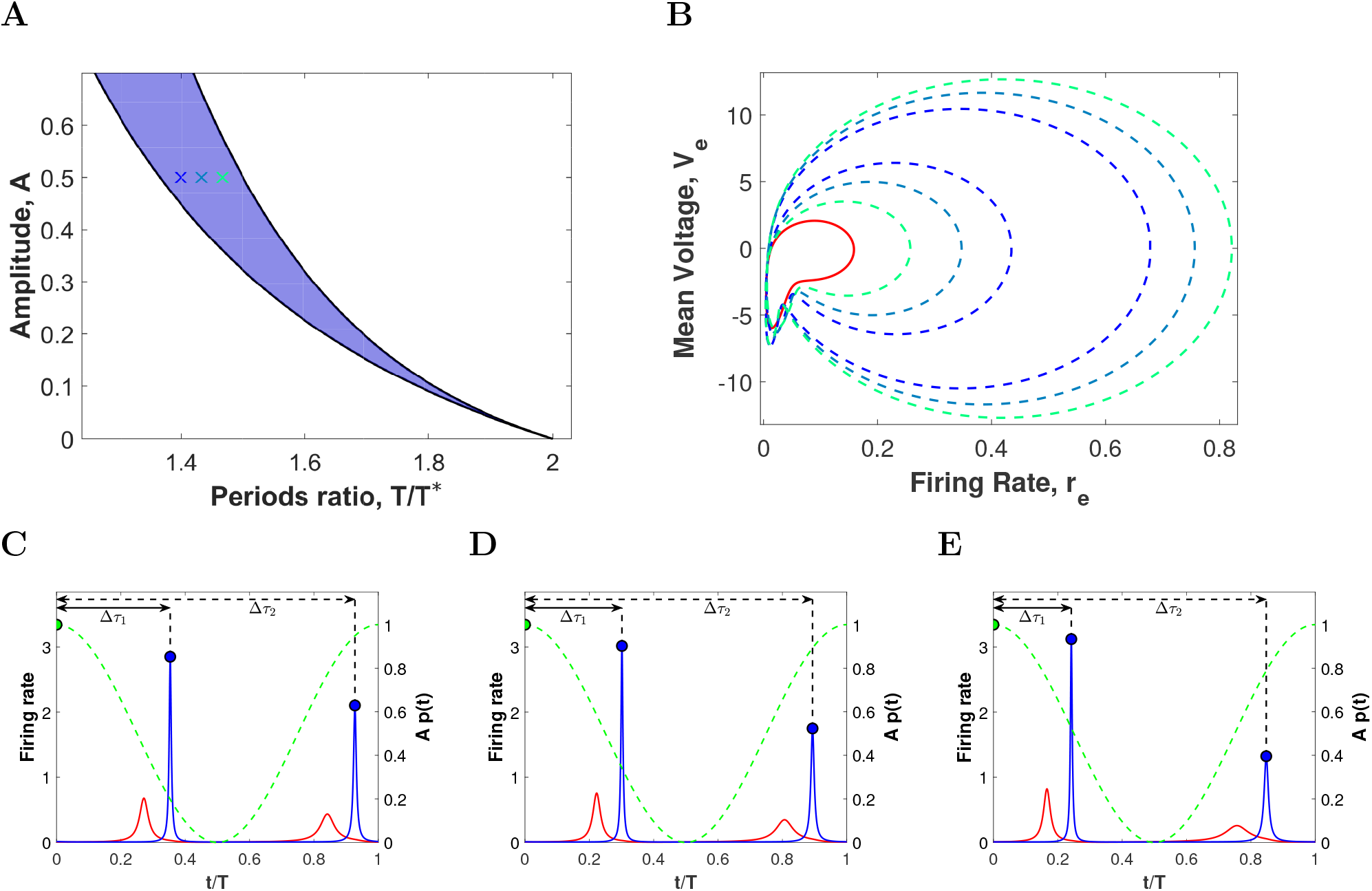
Representative periodic orbits within the 2:1 phase-locking region for sinusoidal inputs. (A) 2:1 phase-locking region between the PING oscillation in Figure 3A and a sinusoidal input (15) varying amplitude factor *A* and period ratio *T/T* ^*^. (B) Projection onto the (*r_e_*, *V_e_*)-plane of 3 representative periodic orbits corresponding to the crosses in panel *A* (dashed curves with same color as the crosses). Solid red line corresponds to the projection of the unperturbed limit cycle. (C-E) Evolution of the firing rate variables *r_e_* (red) and *r_i_* (blue) and the sinusoidal input *Ap*(*t*) (green dashed line) over a single period *T* for the three representative orbits: (C) left-hand side of the Arnold tongue (blue cross), (D) center of the Arnold tongue (turquoise cross) and (C) right hand side of the Arnold tongue (green cross). Blue and green circles indicate the maximum of the I-cells firing rate and the input, respectively. The magnitude Δ*τ*_1,2_ have been pointed out with double-head arrows.

**Figure S5:**
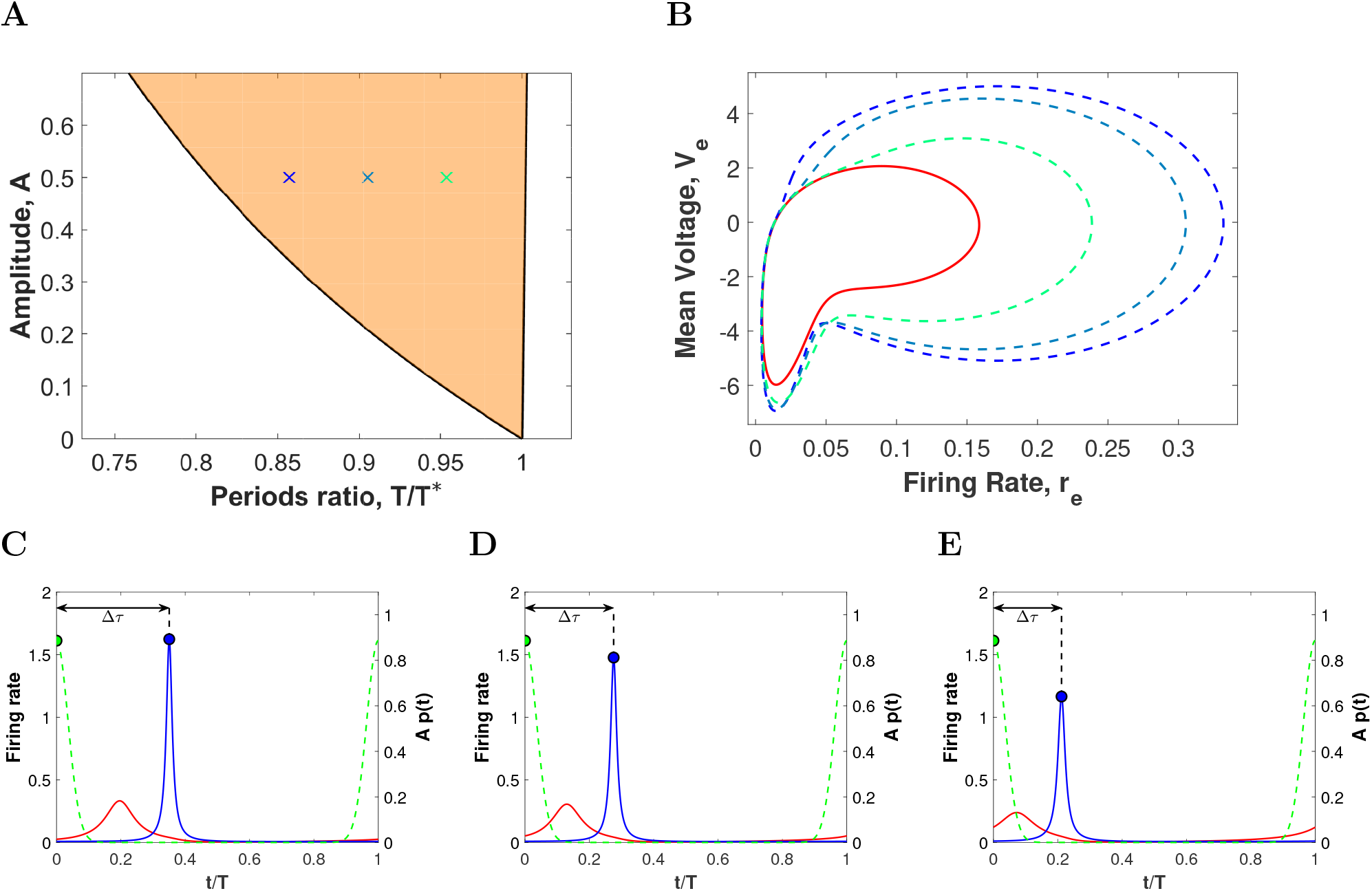
Representative periodic orbits within the 1:1 phase-locking region for high coherence inputs. (A) 1:1 phase-locking region between the PING oscillation in Figure 3A and a von Mises input (17) with *κ* = 20 and varying amplitude factor *A* and period ratio *T/T* ^*^. (B) Projection onto the (*r_e_*, *V_e_*)-plane of 3 representative periodic orbits corresponding to the crosses in panel *A* (dashed curves with same color as the crosses). Solid red line corresponds to the projection of the unperturbed limit cycle. (C-E) Evolution of the firing rate variables *r_e_* (red) and *r_i_* (blue) and the von Mises input *Ap*(*t*) (green dashed line) over a single period *T* for the three representative orbits: (C) left-hand side of the Arnold tongue (blue cross), (D) center of the Arnold tongue (turquoise cross) and (C) right hand side of the Arnold tongue (green cross). Blue and green circles indicate the maximum of the I-cells firing rate and the input, respectively. The magnitude Δ*τ* has been pointed out with a double-head arrow.

**Figure S6:**
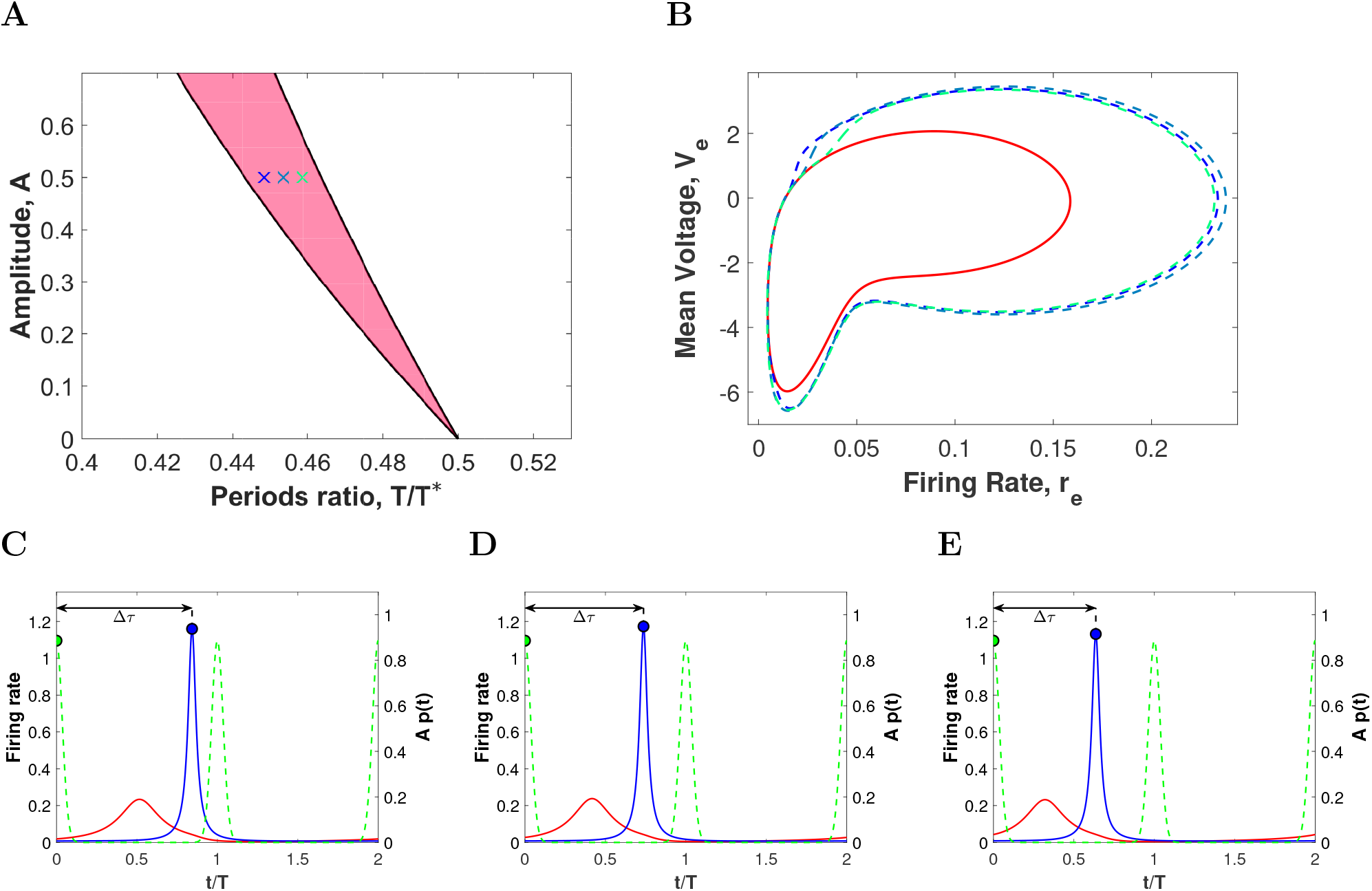
Representative periodic orbits within the 1:2 phase-locking region for high coherence inputs. (A) 1:2 phase-locking region between the PING oscillation in Figure 3A and a von Mises input (17) with *κ* = 20 and varying amplitude factor *A* and period ratio *T/T* ^*^. (B) Projection onto the (*r_e_*, *V_e_*)-plane of 3 representative periodic orbits corresponding to the crosses in panel *A* (dashed curves with same color as the crosses). Solid red line corresponds to the projection of the unperturbed limit cycle. (C-E) Evolution of the firing rate variables *r_e_* (red) and *r_i_* (blue) and the von Mises input *Ap*(*t*) (green dashed line) over two periods *T* for the three representative orbits: (C) left-hand side of the Arnold tongue (blue cross), (D) center of the Arnold tongue (turquoise cross) and (C) right hand side of the Arnold tongue (green cross). Blue and green circles indicate the maximum of the I-cells firing rate and the preceding input volley, respectively. The magnitude Δ*τ* has been pointed out with a double-head arrow.

**Figure S7:**
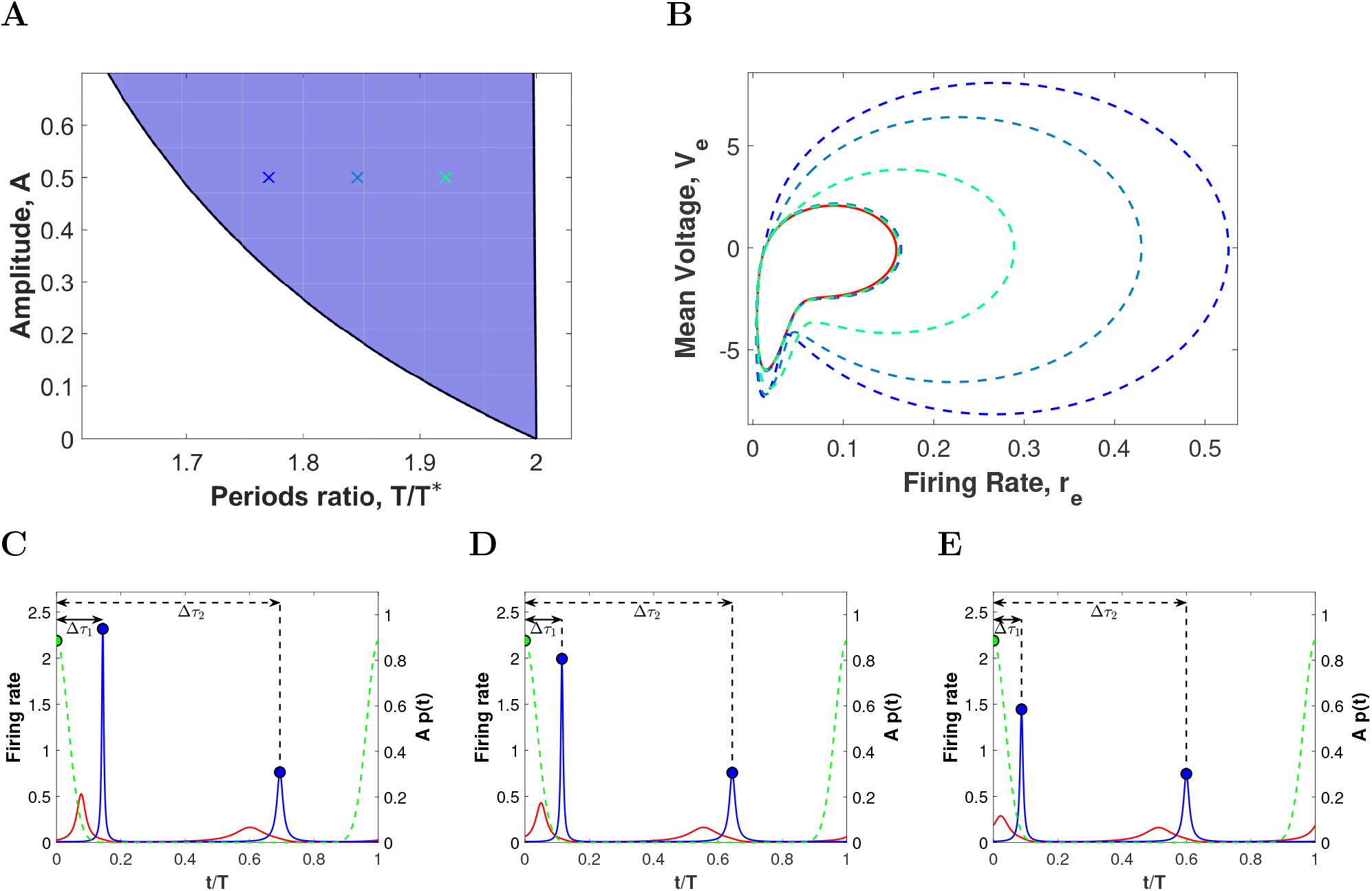
Representative periodic orbits within the 2:1 phase-locking region for high coherence inputs. (A) 2:1 phase-locking region between the PING oscillation in Figure 3A and a von Mises input (17) with *κ* = 20 and varying amplitude factor *A* and period ratio *T/T* ^*^. (B) Projection onto the (*r_e_*, *V_e_*)-plane of 3 representative periodic orbits corresponding to the crosses in panel *A* (dashed curves with same color as the crosses). Solid red line corresponds to the projection of the unperturbed limit cycle. (C-E) Evolution of the firing rate variables *r_e_* (red) and *r_i_* (blue) and the von Mises input *Ap*(*t*) (green dashed line) over a single period *T* for the three representative orbits: (C) left-hand side of the Arnold tongue (blue cross), (D) center of the Arnold tongue (turquoise cross) and (C) right hand side of the Arnold tongue (green cross). Blue and green circles indicate the maximum of the I-cells firing rate and the input, respectively. The magnitudes Δ*τ*_1,2_ have been pointed out with double-head arrows.

**Figure S8:**
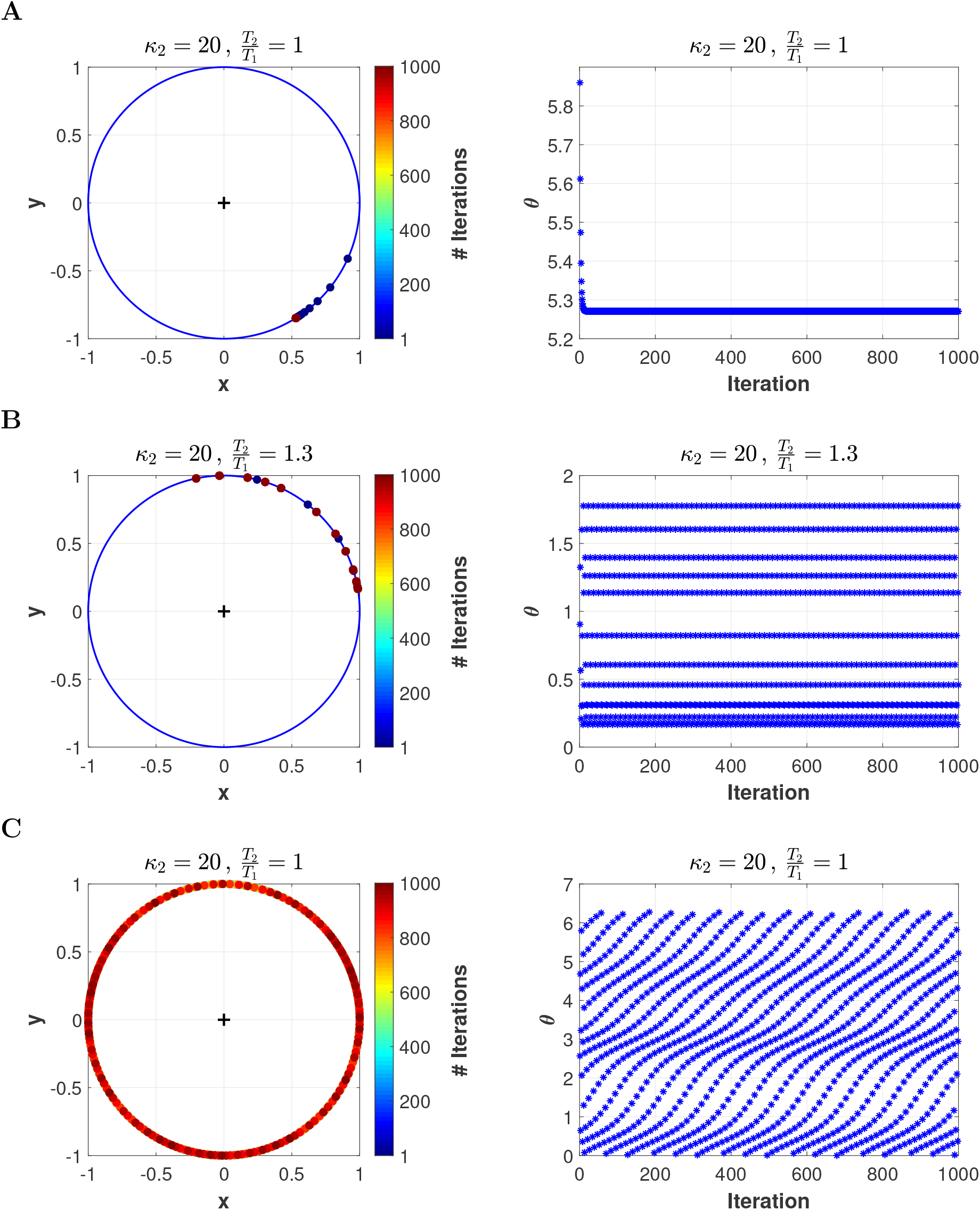
Phase distribution along the unit circle and synchronization index. Phases of the (non-autonomous) stroboscopic map (at time *T*_1_) with two periodic inputs (primary and distractor) of von Mises type with the same coherence value *κ*_1_ = *κ*_2_ = 20 and amplitude *A*_1_ = *A*_2_ = 1. The period varies: (A) *T*_1_/*T* ^*^ = 0.71 and *T*_2_ = *T*_1_, (B) *T*_1_/*T* ^*^ = 0.85 and *T*_2_ = 1.3*T*_1_ and (C) *T*_1_/*T* ^*^ = 1 and *T*_2_ = *T*_1_. (Left column) Distribution of the first 1000 phases along the unit circle and (Right column) plot of the same phases (within the range [0, 2*π*]) as a function of the iterates. The values of the synchronization index for each case are: (A) 0.999, (B) 0.851 and (C) 0.032 (see Figure 8). Three different outcomes are observed in the way the phases are distributed: (A) the iterates converge to a single phase value, (B) the iterates jump from one phase to another within a set of different phase values and (C) the phases fill densely the circle.

**Figure S9:**
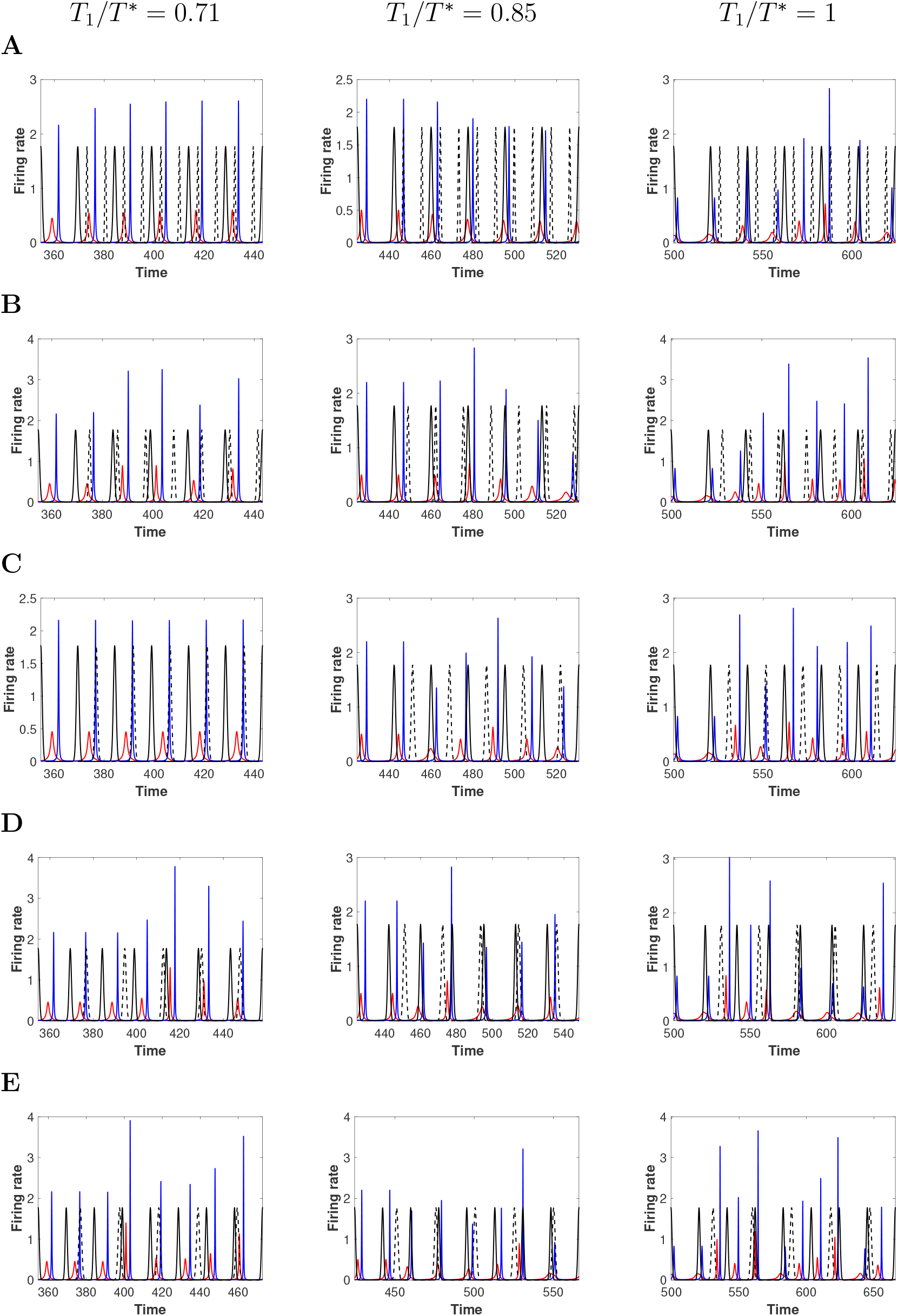
Simulations of the mean field model receiving a primary input and a distractor. Firing rates *r_e_* (red) and *r_i_* (blue) of the mean field model (1)-(2) before and after applying a distractor to the entrained oscillator by the primary input. The primary input of von Mises type (solid black line) oscillates with a relative frequency of *T*_1_/*T* ^*^ = 0.71 (left column), *T*_1_/*T* ^*^ = 0.85 (middle column) and *T*_1_/*T* ^*^ = 1 (right column) with respect to the unperturbed oscillator and it comes in the form of coherent pulses (*κ*_1_ = 20). The distractor is chosen to be as coherent as the first input (*κ*_2_ = 20). The frequency relationship between the primary stimulus and the distractor is (A) *T*_2_/*T*_1_ = 0.5, (B) 0.75, (C) 1, (D) 1.2 and (E) 1.4. The parameters have been chosen as in Figure 8 so that the plots herein illustrate the corresponding predictions. The simulations consist in the integration of system (1)-(2) with the primary stimulus for 25*T*_1_ cycles (only the last cycle is displayed). After that, the system is additionally perturbed with a distractor over a span of 10 max(*T*_1_, *T*_2_) (only the first 5 cycles are shown).

**Figure S10:**
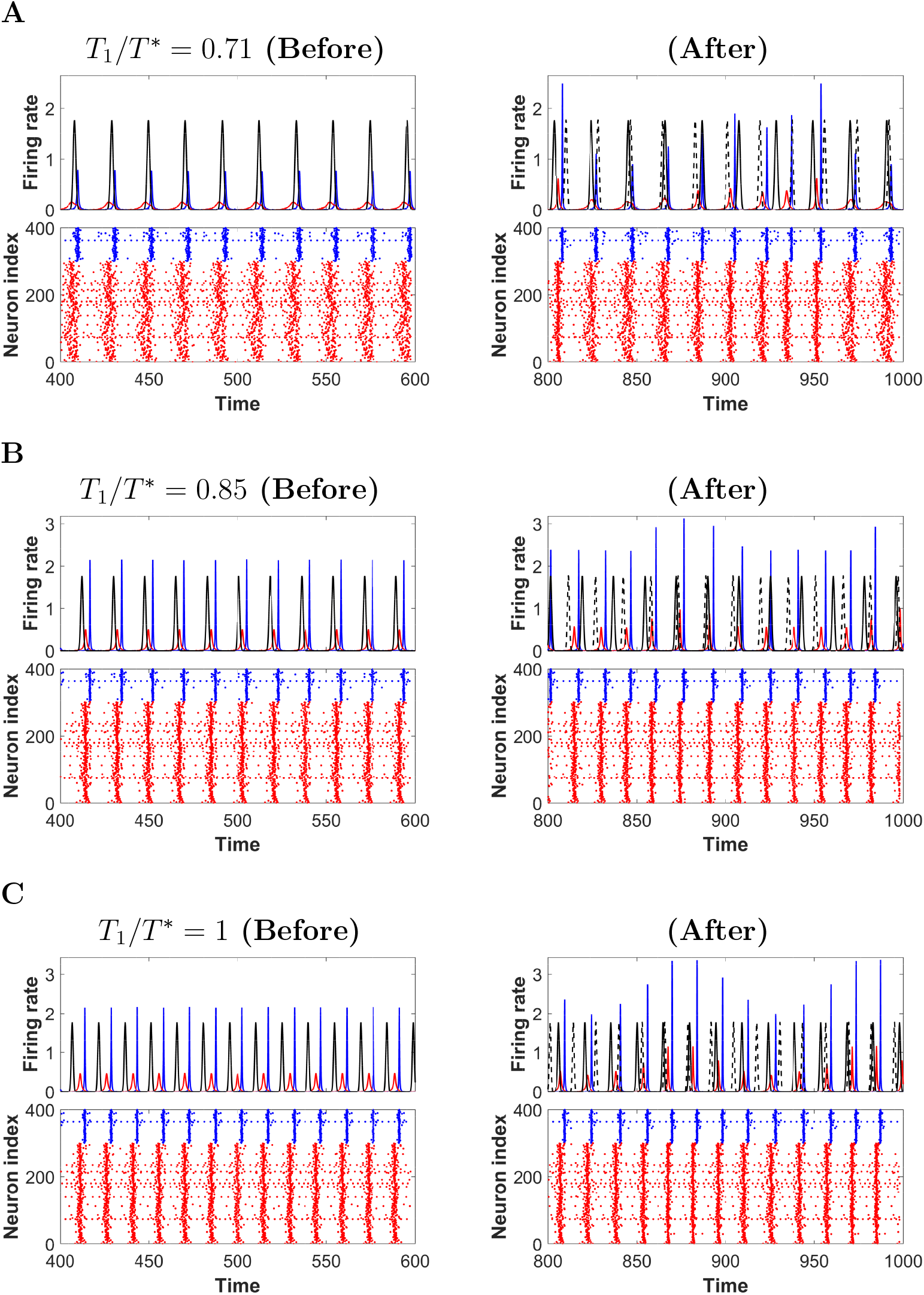
Simulations of the full spiking QIF model receiving a primary input and a distractor. Both inputs are of von Mises type with the same coherence (*κ*_1_ = *κ*_2_ = 20) and relative frequency *T*_2_/*T*_1_ = 0.875. The parameters for the primary have been chosen within the 1:1 Arnold tongue as in Figure 8: *A*_1_ = 1 and (A) *T*_1_/*T* ^*^ = 0.71 (left hand side of the Arnold tongue), (B) *T*_1_/*T* ^*^ = 0.85 (middle of the Arnold tongue) and (C) *T*_1_/*T* ^*^ = 1 (right hand side of the Arnold tongue). For each plot we show the mean firing rates and raster plots before (left column) and after (right column) applying the distractor at time 600 ms (each panel shows 200 ms of integration time).

